# Leveraging a natural murine meiotic drive to suppress invasive populations

**DOI:** 10.1101/2022.05.31.494104

**Authors:** Luke Gierus, Aysegul Birand, Mark D. Bunting, Gelshan I. Godahewa, Sandra G. Piltz, Kevin P. Oh, Antoinette J. Piaggio, David W. Threadgill, John Godwin, Owain Edwards, Phillip Cassey, Joshua V. Ross, Thomas A. A. Prowse, Paul Q. Thomas

## Abstract

Invasive rodents, including house mice, are a major cause of environmental damage and biodiversity loss, particularly in island ecosystems. Eradication can be achieved through the distribution of rodenticide, but this approach is expensive to apply at scale, can have negative impacts (e.g. on non-target species, or through contamination), has animal ethics concerns, and has restrictions on where it can be used. Gene drives, which exhibit biased inheritance, have been proposed as a next generation strategy to control invasive alien pests and disease vectors. However, synthetic gene drives including CRISPR homing drives have proven to be technically challenging to develop in mice. The *t* haplotype is a naturally-occurring segregation distortion locus with highly biased transmission from heterozygous males. Here we propose a novel gene drive strategy for population suppression, *t*_CRISPR_, that leverages *t* haplotype bias and an embedded SpCas9/gRNA transgene to spread inactivating mutations in a haplosufficient female fertility gene. Using spatially explicit individual-based *in silico* modelling, we show that polyandry, sperm competition, dispersal, and transmission bias are critical factors for *t*_CRISPR_-mediated population suppression. Modelling of realistic parameter values indicates that *t*_CRISPR_ can eradicate an island population of 200,000 mice while the unmodified *t* haplotype fails under the same conditions. We also demonstrate feasibility of this approach by engineering *t*_CRISPR_ mice in a safe split drive format. *t*_CRISPR_ mice exhibit high transmission of the modified *t* haplotype, and efficient generation and transmission of inactivating mutations in a recessive female fertility gene, crucially, at levels for which the modelling predicts that population eradication can occur. This is the first example of a feasible gene drive system for invasive alien rodent population control.

## Introduction

Invasive mammalian pests are among the greatest threats to global biodiversity and have constituted an unprecedented form of global change^1,2^. Commensal rodents, including house mice (*Mus musculus*), have spread throughout the globe, causing significant environmental damage and loss of agricultural productivity^3,4^. Islands are biodiversity hotspots and are particularly susceptible to the impact of invasive rodents, where they contribute to widespread extinction and endangerment, particularly of migratory bird species, reptiles, and plant stocks that have not evolved with rodents^5–9^. Current control methods rely principally on widespread distribution of anticoagulant toxicants, an approach that is not species specific, only feasible on islands without significant human population or livestock, costly to apply at a scale, can have negative impacts (e.g. on non-target species, or though contamination) and has animal ethics concerns^10^.

Genetic biocontrol technologies including gene drives offer the potential for landscape-scale modification or suppression of invasive populations^11^. Gene drives are natural or synthetic genetic elements that spread through a population via super-Mendelian transmission. Building upon the pioneering concepts of Austin Burt^12^ and others^13^, and leveraging the emergence of CRISPR/Cas9 gene editing technology^14^, synthetic “homing” gene drives have recently been developed in several insect species, including *Anopheles* malaria vectors^15–18^. However, in mice, homing gene drives have been challenging to develop^19–21^, prompting us to consider alternative strategies. First described in 1927^22^, the *t* haplotype is a naturally-occurring gene drive element that is commonly found in wild mouse populations that functions as a male meiotic drive, biasing transmission from carrier males by up to ~95%, depending on the variant^23,24^. In non-lethal *t* haplotype variants such as *t*^w2^, homozygous males are sterile and homozygous females are viable and fertile^25^. Intriguingly, despite the high transmission of the *t* haplotype, the frequency within mainland populations remains relatively low (13-55% in populations with predominantly non-lethal *t* haplotype^26^, a phenomenon known as the ‘*t* paradox’^27^). This effect appears to be due to polyandry (i.e. mating with multiple males per breeding cycle) and sperm competition, which forces subfunctional *t* sperm to compete with fully fit sperm from wild type males^28–31^. However, non-lethal *t* haplotypes have not been found in island populations (apart from one report^32^). Given the significant bias in *t* transmission, it is possible that this natural male meiotic drive could be leveraged for mouse population suppression or even eradication on islands^3^. Here we describe a novel gene drive strategy termed *t*_CRISPR_, in which fertile females are progressively depleted due to a CRISPR transgene embedded in the *t* haplotype that targets a haplosufficient female fertility gene. Using spatially explicit individual-based *in silico* modelling on a hypothetical island, we demonstrate that *t*_CRISPR_ has eradication potential across a range of realistic scenarios. We further demonstrate the feasibility of this strategy by engineering and testing *t*_CRISPR_ in a genetically-contained “split drive” format, where the Cas9 and gRNA expression cassettes are integrated into different chromosomes.

## Results

### *In silico* spatial modelling demonstrates eradication potential of *t*_CRISPR_

Given the significant transmission bias of the *t* haplotype, we investigated whether this could be harnessed to develop a synthetic CRISPR gene drive targeting a haplosufficient gene required for female fertility (e.g. Prolactin, *Prl*). We envisaged a system whereby mutations in *Prl* are generated in the male germline through activation of a Cas9/gRNA transgene embedded in a non-lethal *t* variant such as *t*^w2^ (which is transmitted to ~95% of offspring^33^ by heterozygous males (*t*_CRISPR_; Fig. 1a,b). To assess the population suppression potential of the *t*_CRISPR_ strategy, we performed spatially explicit individual-based simulations, which demonstrated that *t*_CRISPR_ can effectively eradicate large populations (*N* ~ 200,000) of mice (Figs. 2a, 3). Typically, during the first decade after inoculation, *t*_CRISPR_ spread across the entire landscape, *Prl* (functional) allele frequency declined steadily, and substantial (> 90%) suppression of the population was achieved in ~20 years (e.g. Fig. 2a), with complete eradication in ~28 years. Simulated eradication using *t*_CRISPR_ was successful across a wide range of parameters related to gene drive efficiency and mouse life history, often with shorter time to eradication (see below). *t*^w2^ can also effect population suppression through sterility of male homozygotes (e.g. Fig. 2b); however, simulated eradication probability was much lower compared to *t*_CRISPR_, and when eradication was successful, the time to eradication was substantially longer (Fig. 3). Time to eradication for *t*_CRISPR_ was slightly longer than other proposed gene-drive strategies such as a homing drive for female infertility and a “driving Y” X-shredder^30–32^ (Extended Data Fig. 1).

**Figure 1.**
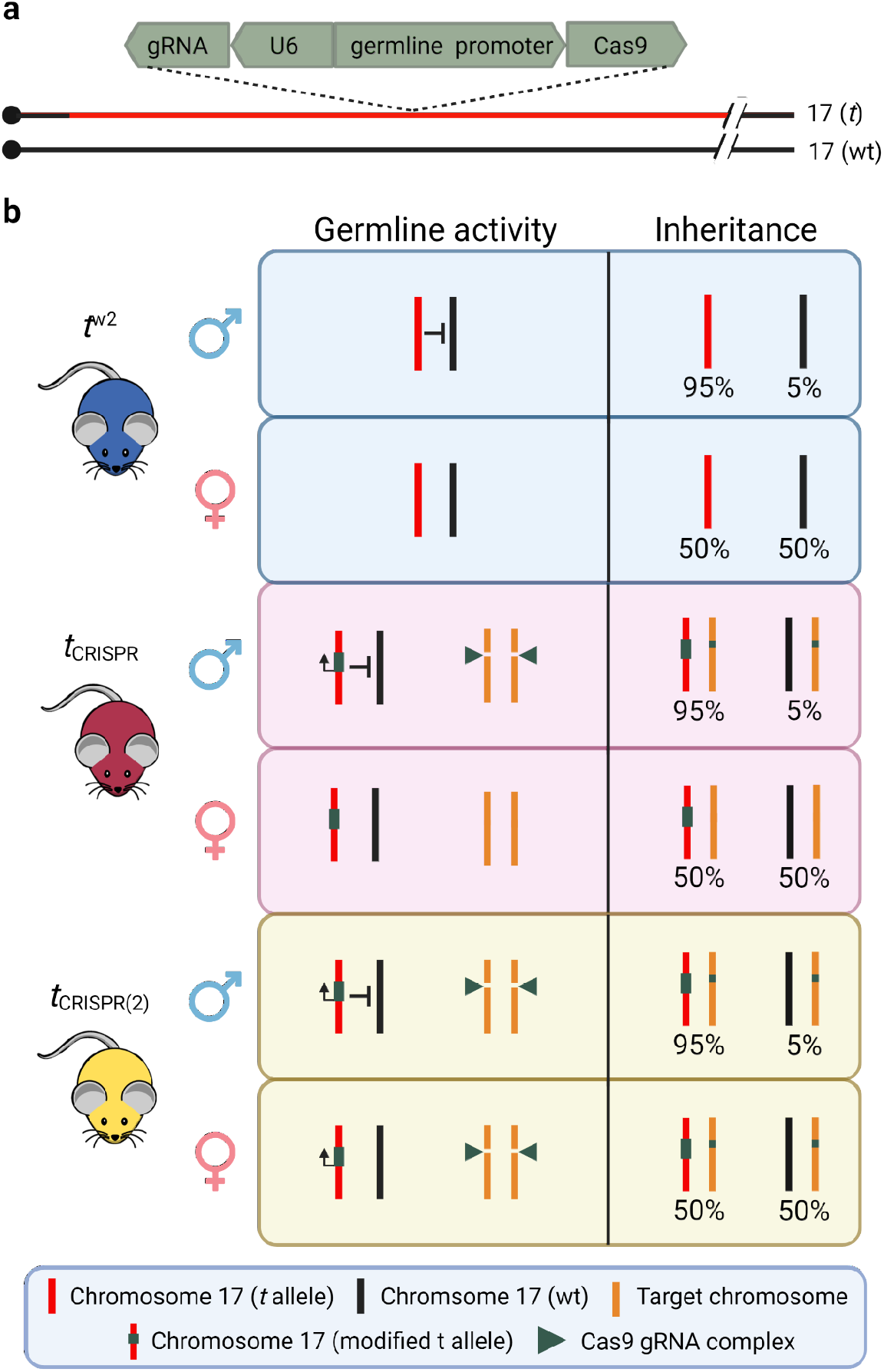
Overview of *t* haplotype modification strategies for population suppression. **a.** The integration of a transgene within the *t* haplotype expressing Cas9 under the control of a male specific promoter or a germline specific promoter, coupled with a ubiquitously-expressed gRNA targeting a haplosufficient female fertility gene. **b.** Inheritance of *t*^w2^ is biased in males but not females. In the *t*_CRISPR_ system, Cas9 is only active in males and, with a ubiquitously expressed gRNA, disrupts a haplosufficient female fertility gene in the germline. *t*_CRISPR_ males transmit the *t*_CRISPR_ transgene and a disrupted fertility gene to 95% and 100% of offspring, respectively. The *t*_CRISPR(2)_ strategy is identical except that the Cas9 is active in the male and female germline, thereby generating a more rapid increase in female infertility alleles.

**Figure 2.**
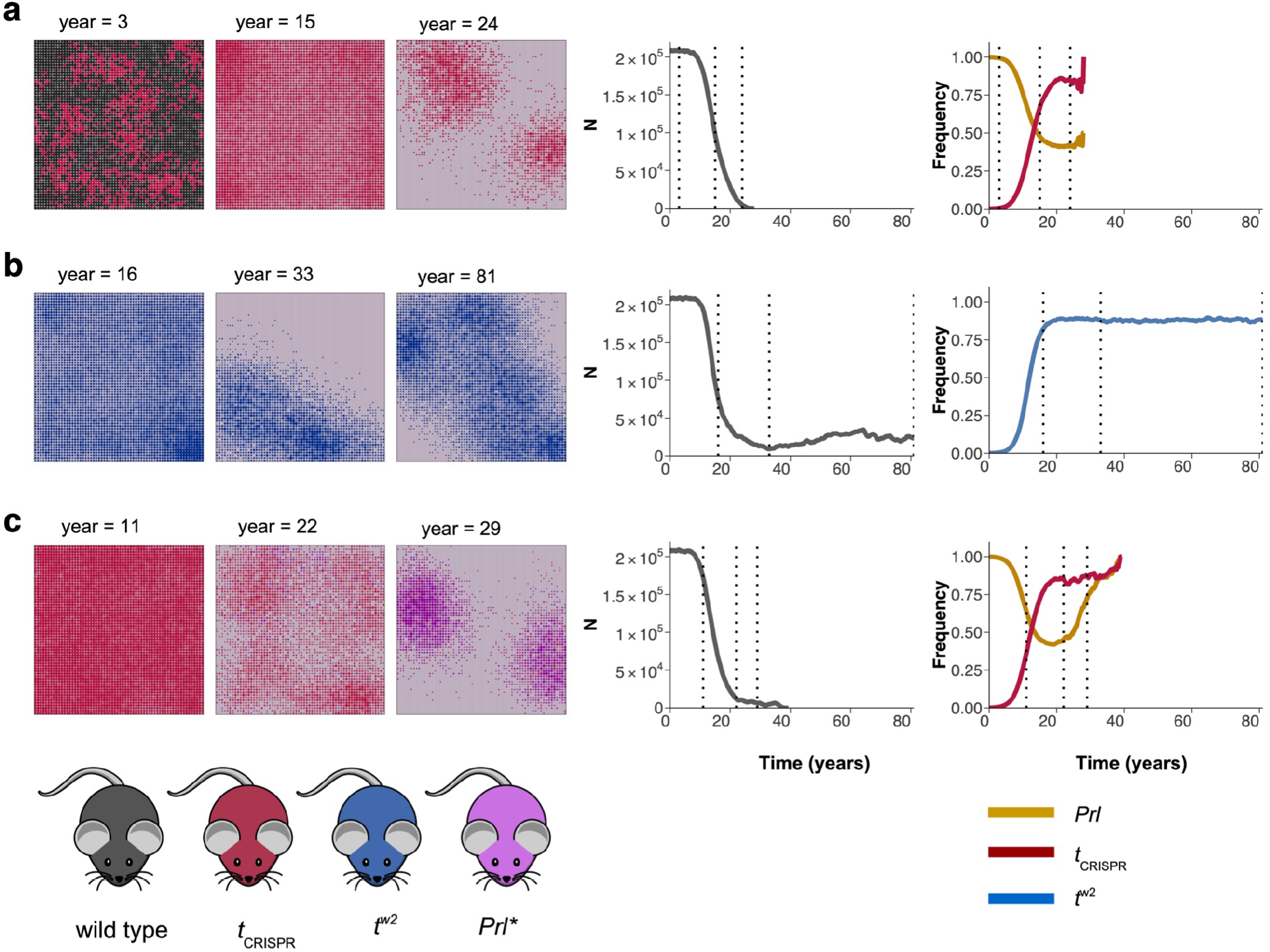
Sample simulations using two strategies, *t*_CRISPR_ and *t*^w2^. Shown in order are three snapshots of the spatial distribution of mouse populations, total population size and the allele frequency change through time, starting with the inoculation of *t* haplotype individuals at year = 0 (dashed lines in the plots correspond to the years of the snapshots). Each circle represents the population in that patch: the circle size is proportional to the patch population size, and the circle color represents the most abundant *t* haplotype in that patch. Empty patches appear as light gray. (See Extended Data Table 1 for the base values of the parameters used, and Supplementary Materials Online for higher resolution images of the snapshots and for videos of the simulations.) **a.** *t*_CRISPR_ spread across the entire landscape in ~10 years and substantial suppression of the population was achieved after 21 years with complete eradication in 28.1 years. Functional prolactin frequency declined steadily in the population following inoculation with *t*_CRISPR_ individuals. **b.** *t*^w2^ spread across the entire landscape in similar time frames to *t*_CRISPR_, but eradication of the population was not achieved. **c.** When the probability of loss of function after a successful DNA cut is reduced to *p*_L_ = 0.9996, resistant genotypes (*Prl*–) emerged in ~20 years after inoculation and functional prolactin levels started to increase, but eradication was still successful (*p*_m_ = 0.63 in all three simulations).

**Figure 3.**
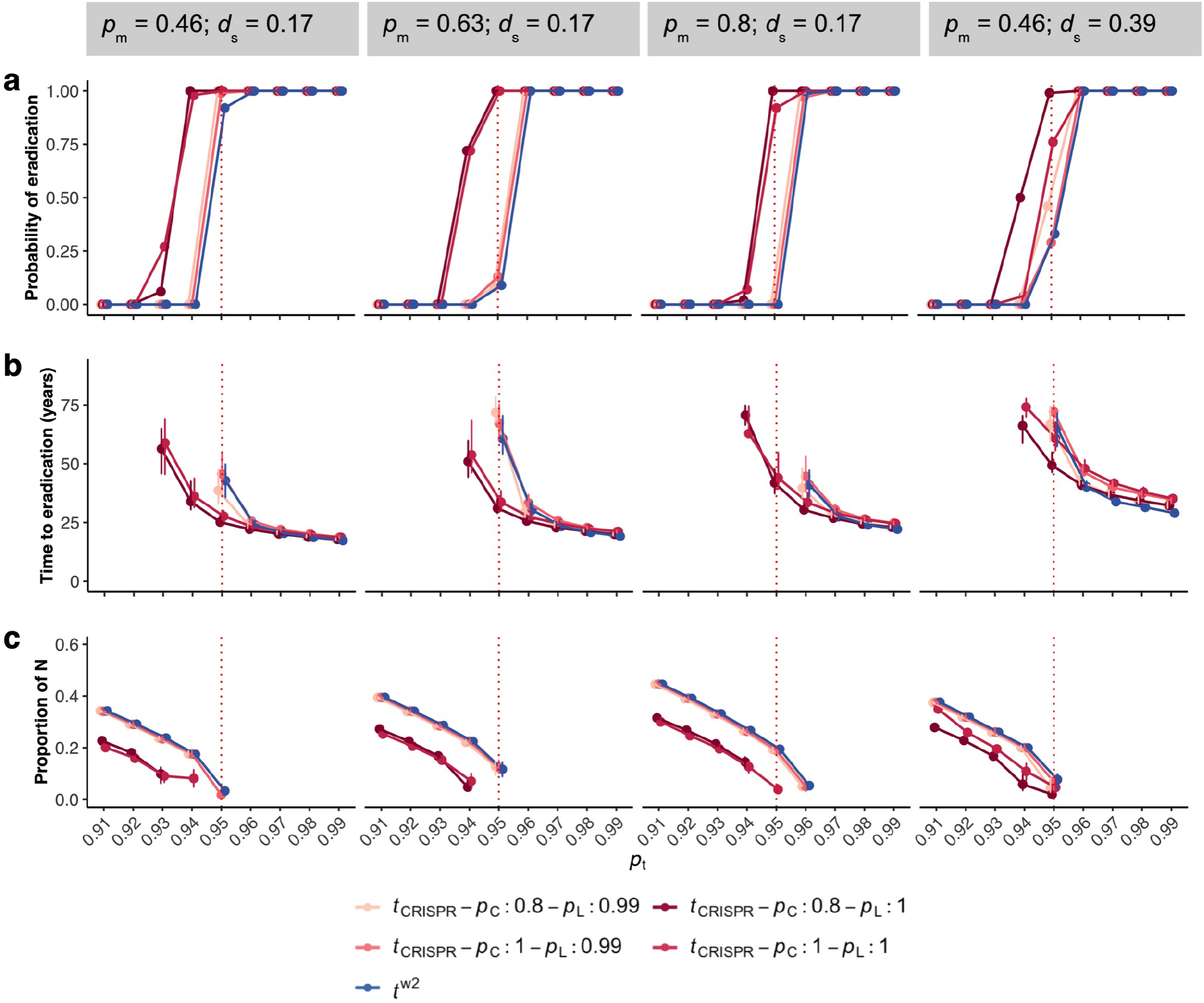
Effect of probability of transmission *p*_t_ on the success of the *t*^w2^ and *t*_CRISPR_ elimination strategies. **a**. The probability of eradication; the median with interquartile ranges of the time to eradication (**b**), and the proportion of remaining population sizes when eradication was unsuccessful (**c**) based on 100 simulations for each parameter combination (18,000 simulations in total). Higher values of *p*_t_ lead to higher probabilities of eradication, shorter times to eradication, and higher population suppression when eradication failed, whereas high levels of either sperm competition (*d*_s_ = 0.39) or polyandry (*p*_m_ = 0.8) reduced eradication probabilities and delayed times to eradication except when *p*_t_ was very high (**b**). Eradication failed for both strategies when sperm competition and polyandry are very high (results not shown when *p*_m_ ≥ 0.63 and *d*_s_ = 0.39). Compared to *t^w2^*, *t*_CRISPR_ generally had higher eradication probabilities, shorter eradication times, and higher population suppression when eradication failed. At *p*_t_ = 0.95 (dashed line), *t*_CRISPR_ was always successful if there is complete loss of function (*p*_L_ = 1).

The transmission probability of the *t* haplotype in heterozygous males (*p*_t_) and probability of polyandry (*p*_m_) strongly influenced *t*_CRISPR_ eradication probability (Fig. 3a) and time to eradication (Fig. 3b). Eradication is successful when *p*_t_ ≥ 0.93 with moderate levels of polyandry (*p*_m_ = 0.46). Higher levels of polyandry or sperm competition require higher transmission efficiencies (*p*_t_) for successful eradication. With these parameter values, *t*^w2^ produced very low probabilities of eradication. In contrast, when *p*_t_ = 0.95, *t*_CRISPR_ successfully eradicated at all levels of polyandry tested (*p*_m_ = 0.46, 0.63, 0.8) when sperm competition was moderate (*d_s_* = 0.17), and at moderate levels of polyandry when sperm competition was very high (*d*s = 0.39). In addition, time to eradication was much shorter with *t*_CRISPR_ than with *t*^w2^ (~25 years and ~43 years for *t*_CRISPR_ and *t*^w2^, respectively, when *p*_m_ = 0.46, *d*_s_ = 0.17; Fig. 3b). *t*_CRISPR_ also achieved stronger population suppression than *t*^w2^ when eradication failed for both strategies (Fig. 3c).

The probability of a successful DNA cut (*p*_C_) had an unexpected effect on simulation outcomes where less efficient cutting (e.g. 0.8) resulted in a higher probability of eradication and shorter time to eradication than *p*_C_ = 1.0 (Fig. 3, Extended Data Fig. 2). In contrast, a high probability of loss of gene function (*p*_L_) following a successful cut is required to prevent resistant alleles that retain function from emerging (e.g. Fig. 2c with *p*_C_ = 0.8 and *p*_L_ = 0.9996). Typically, once functional resistant genotypes emerged, the decline in *Prl* frequency was rapidly reversed.

However, in most cases eradication was still achieved with *t*_CRISPR_ when *p*_L_ < 1 (Extended Data Fig. 2). *t*_CRISPR_ continued to outperform *t*^w2^ for all values of *p*_C_ and *p*_L_ when polyandry and sperm competition were moderate, and when *p*_C_ < 0.95 for higher values of polyandry and sperm competition. This suggests that eradication was achieved mostly through male sterility; however, female infertility continued to contribute to eradication (Extended Data Fig. 2). When *p*_L_ was reduced even further to 0.99, the contribution of female infertility declined and *t*_CRISPR_ behaved more like *t*^w2^ with a similar probability of eradication and time to eradication (Fig. 3).

The predictions from the Boosted Regression Tree model, fitted to the sensitivity-analysis results, support the above predictions (Fig. 4, Extended Data Fig. 3). *t*_CRISPR_ was predicted to successfully eradicate for a wide range of transmission probabilities (*p*_t_ ≥ 0.93) and polyandry (*p*_m_ > 0.2) when sperm competition was moderate (*d_s_* = 0.17, Fig. 4a; also see Extended Data Fig. 4a). When *p*_t_ = 0.95, *t*_CRISPR_ was predicted to successfully eradicate and could tolerate higher levels of both sperm competition and polyandry, whereas *t*^w2^ was successful only for low levels of polyandry (*p*_m_ < 0.4, Fig. 4b, Extended Data Fig. 4b). Increasing dispersal distance (*D*) increased eradication probabilities for *t*_CRISPR_ when *p*_t_ < 0.96, but not for *t*^w2^ (Fig. 4c, Extended Data Fig. 4c). Lower transmission probabilities *p*_t_, higher levels of polyandry and sperm competition delayed the time to eradication for both strategies (Fig. 5); however, *t*_CRISPR_ had shorter time to eradication than *t*^w2^ for all values of polyandry and sperm competition when *p*_t_ ≤ 0.95 (also see Extended Data Fig. 5).

**Figure 4.**
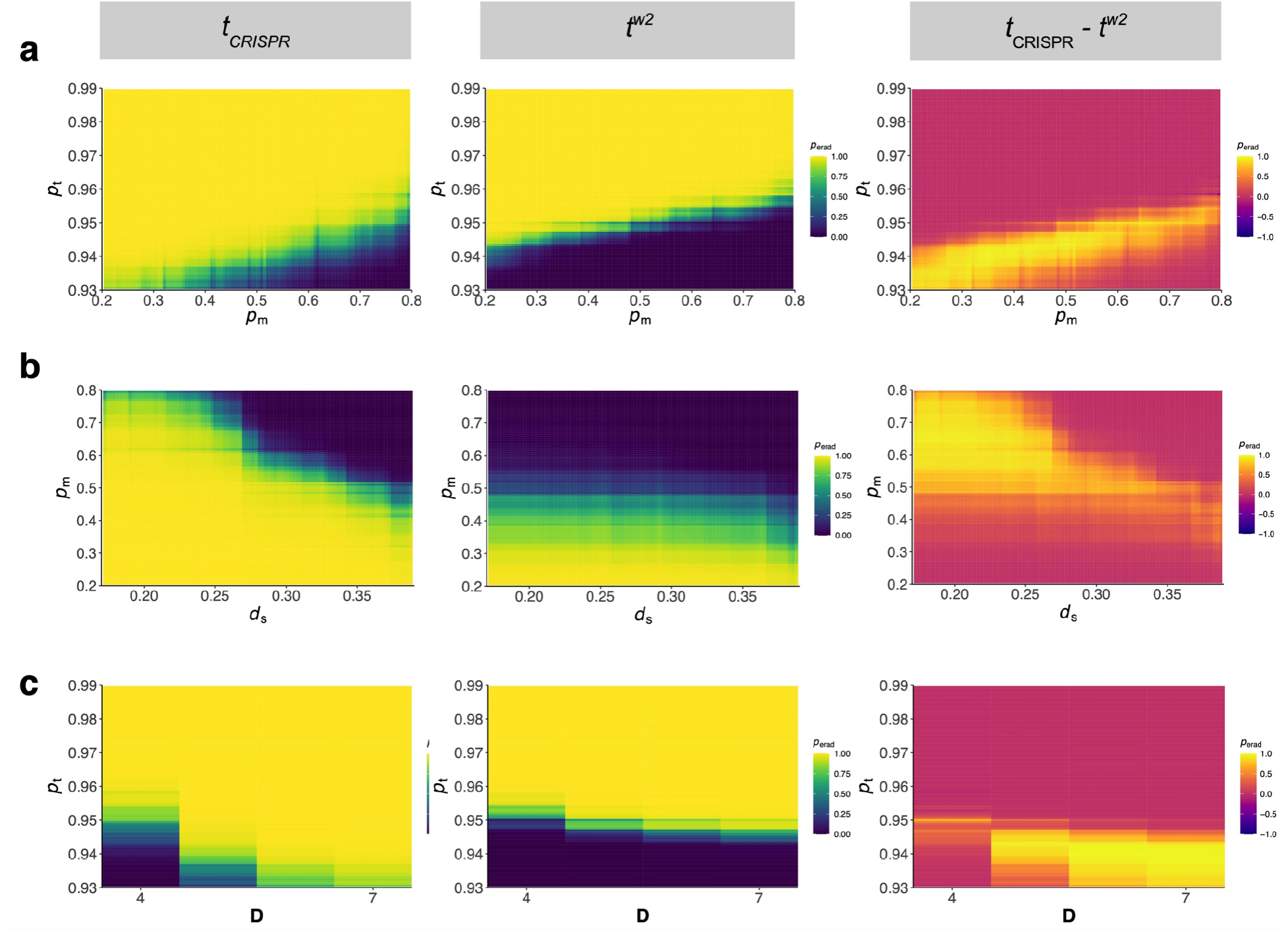
Predicted probabilities of eradication based on Boosted Regression Tree model fitted to sensitivity-analysis results. Shown are the probabilities of eradication using *t*_CRISPR_ (left) and *t*^w2^ (middle), and their difference (right; note the scale change to [-1, 1]). **a.** *t*_CRISPR_ was predicted to be successful in eradication for less efficient transmission (*p*_t_ < 0.95) together with low to moderate levels of polyandry (*p*_m_ < 0.46), where *t*^w2^ failed (assuming moderate sperm competition, *d*_s_ = 0.17). At *p*_t_ = 0.95, *t*_CRISPR_ was predicted to be successful for a wide range of *p*_m_. **b.** The expected probability of eradication was very low with *t*^w2^ for *p*_m_ > 0.40 and zero for *p*_m_ > 0.60 irrespective of levels of sperm competition (assuming *p*_t_ = 0.95). *t*_CRISPR_ had higher probabilities of eradication for higher levels of polyandry, unless sperm competition was very high. **c.** Dispersal distances, *D* ≥ 5 increased eradication probabilities for *t*_CRISPR_ when *p*_t_ ≤ 0.95, but not for *t*^w2^ (assuming *p*_m_ = 0.46). Other parameters, if not stated, are given in Extended Data Table 1.

**Figure 5.**
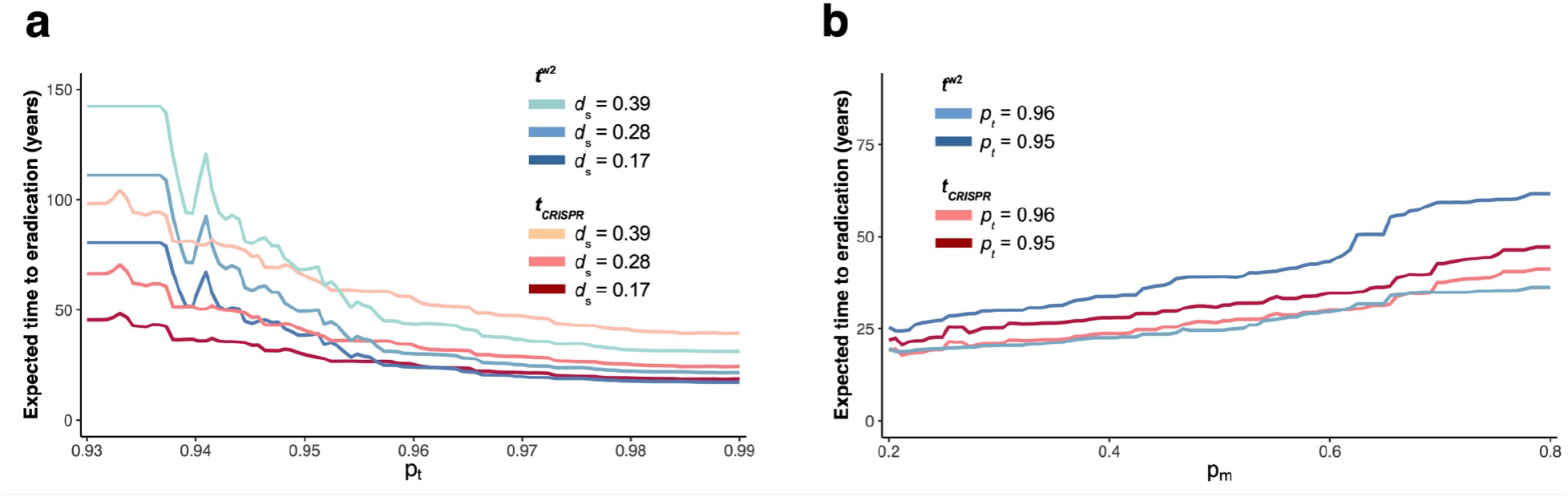
Expected times to eradication. Predictions are based on the Boosted Regression Tree model fitted to sensitivity-analysis results where eradication was successful (1641 simulations using *t*_CRISPR_, and 1634 simulations using *t*^w2^). **a.** Low transmission probabilities *p*_t_, delayed the time to eradication for both strategies; but more so for *t*^w2^. Increasing sperm competition also delayed the time to eradication for both strategies (*p*_m_ = 0.46). **b.** Increasing polyandry *p*_m_, delayed the time to eradication, particularly for *t*^w2^ when *p*_t_ = 0.95 (*d*_s_ = 0.17).

We also investigated eradication probability with a related strategy (*t*_CRISPR(2)_) in which Cas9 is active in the germline of both males and females (Extended Data Fig. 6). Although *Prl* is not required in the female germline, inactivation of *Prl* in females ultimately caused a critical shortage of fertile females carrying the *t* haplotype (Extended Data Fig. 6b), which limited drive spread. Given that the *t* haplotype is present in some wild populations^26^, we also modelled the impact of endogenous *t*^w2^ in the target population on the eradication success of *t*_CRISPR_. Since *Prl* prevalence is reduced after the introduction of the drive, *t*_CRISPR_ was at a disadvantage with fewer fertile females spreading the drive compared to the naturally occurring *t* haplotypes (e.g. Extended Data Fig. 7a). Encouragingly, analysis of previously published whole-genome pool-seq datasets^34^ (*n* = 40 individuals per island) for reads mapping to diagnostic transcripts detected no evidence of *t* haplotypes among mice sampled from four islands in Western Australia, the north Pacific, and the western coast of North America (Extended Data Fig. 8). In contrast, one of the four “continental” populations in the same study showed evidence of *t* haplotypes, with the number of reads mapped exceeding the expected values at the lowest detectable *t* haplotype frequency (i.e., a single *t* haplotype sampled). Interestingly, our results suggest that the competitive advantage of naturally occurring *t* haplotypes over *t*_CRISPR_ could allow them to be used as “rescue drives”, albeit only under a limited range of parameter values (e.g. Extended Data Fig. 7b).

Initial population size on the island did not affect time to eradication when either the population density or the island size were altered (Extended Data Fig. 9), suggesting that the length and the number of breeding cycles in a year, along with dispersal abilities, influence the temporal dynamics of the drive. Lastly, if *t* carrying females preferred wild type males over *t* carrying males, eradication probability reduced and time to eradication increased (Extended Data Fig. 10); however, *t*_CRISPR_ had better eradication success than *t*^w2^. Together, these predictions indicate that the *t*_CRISPR_ strategy has robust eradication potential across a range of realistic parameter values.

### Generation and analysis of a *t*_CRISPR_ split drive system

To assess the feasibility of the *t*_CRISPR_ strategy, we sought to generate and test a *t*_CRISPR_ drive under laboratory conditions. To ensure containment, we employed a split drive design whereby a constitutive gRNA expression cassette was integrated into the *t*^w2^ locus and a male germline Cas9 expression cassette was inserted at a different genomic location. We selected *Prl* for the gRNA target as the function of this haplosufficient female-specific fertility gene is well characterised^35^. Six gRNAs targeting PRL essential amino acids^36–40^ (P2.1-P2.6) were screened for cutting efficiency (*p*_C_) an the probability of generating a loss of function mutation (*p*_L_). gRNA P2.3 was selected for further investigation based on very high cleavage efficiency (*p*_C_ = 0.997) and predicted loss of function (*p*_L_ = 1.0; see Extended Data Fig. 11).

A P2.3 gRNA expression mouse line (*t*^w2-gRNA^) was created by targeted integration of a U6-gRNA-mcherry expression cassette at an intergenic site within inversion 1 of the *t*^w2^ haplotype^41^ (Extended Data Fig. 13a,b). We used the early spermatogenesis-specific promoter *Ccna1*^42^ to drive Cas9 expression in the male germline (Tg^Cas9^; Extended Data Fig. 13c,d). *t*^w2-gRNA/0^;Tg^Cas9/0^ transhemizygous males (modelling *t*_CRISPR_) were mated with wild type female mice to determine transmission of the modified *t* haplotype, *Prl* indel efficiency and the frequency of functional resistant alleles (Extended Data Fig. 13e). Of the 203 F1 embryos sired by transhemizygous males, 142 (*p*_C_ = 0.70) contained indels at the target site (range 0.364 - 0.891 per sire; Table 1), including one sample with biallelic *Prl* mutations, presumably due to Cas9/gRNA carryover. The average number of embryos sired by *t*_CRISPR_ males (7.8 per pregnant female; Table 1) was not significantly different to the average litter size from *t*^w2^ sires (7.0; *p* = .2620). NGS analysis of *t*^w2-gRNA/0^;Tg^Cas9/0^ sperm revealed a similarly high indel efficiency (*p*_C_ = 0.801, range 0.422 – 0.926 based on 52,427 reads from 10 mice; Table 2). When cutting occurred, the predicted frequency of non-functional *Prl* alleles was 1.0. *t* haplotype transmission from *t*^w2-gRNA^ males was 0.951 indicating that the transgene insertion did not alter *t* haplotype function. *t*^w2-gRNA/0^;Tg^Cas9/0^ females did not display biased transmission of the modified *t* haplotype (*p*_t_ = 0.519; Table 1) and no indels were observed in offspring (*n* = 27) demonstrating that Cas9 activity is confined to males, as expected. Together, these data demonstrate the key properties of the *t*_CRISPR_ drive: biased transmission of a modified *t* haplotype and efficient generation and transmission of inactivating mutations in a recessive female fertility gene.

**Table 1:**
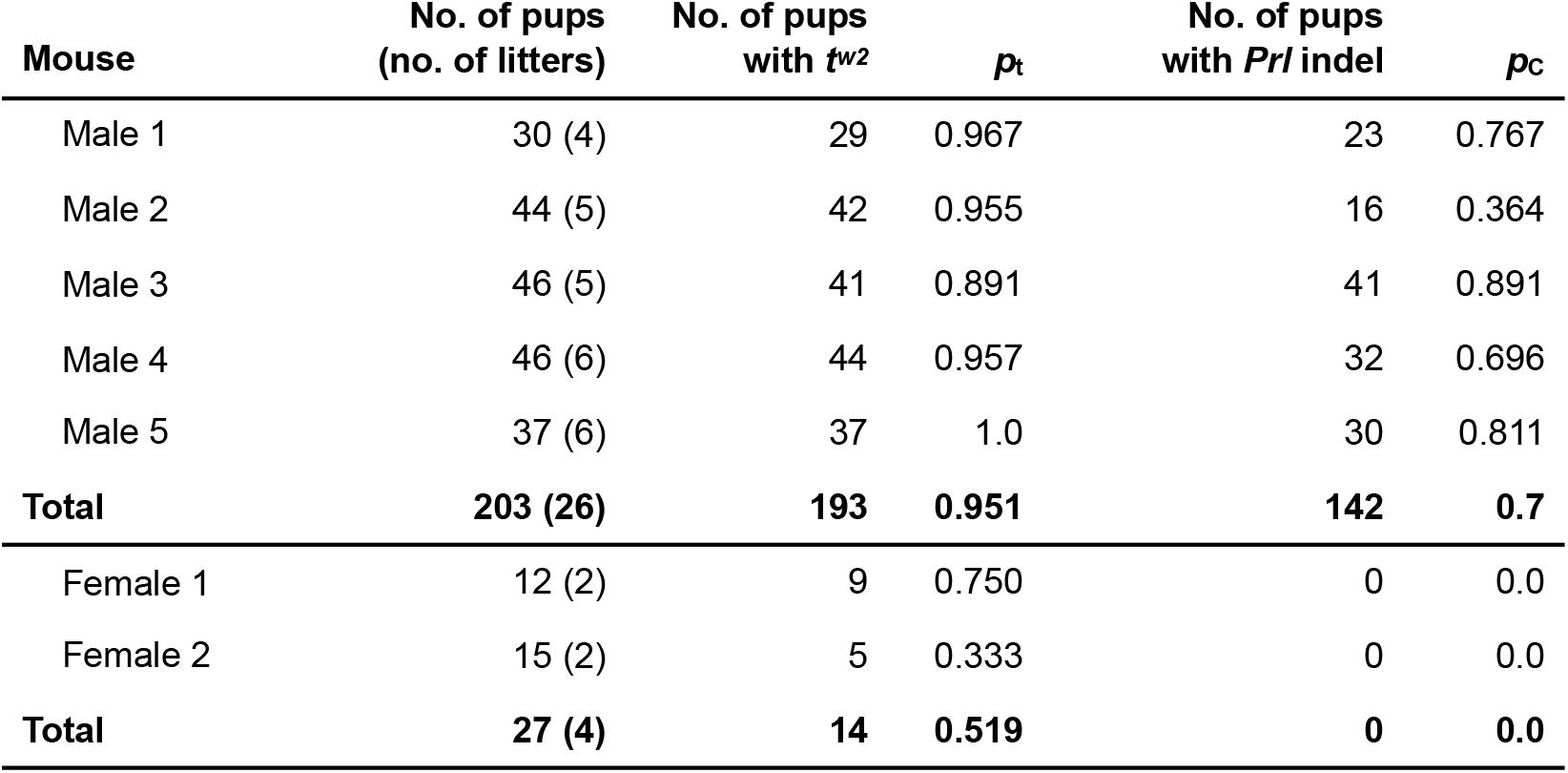
Modified *t*^w2^ haplotype transmission, *p*_t_, of transhemizygous *t*_CRISPR_ mice mated with wild type C57BL/6 mice. Pups were screened for P2.3 gRNA cutting efficiency, *p*_C_.

| Mouse | No. of pups<br>(no. of litters) | No. of pups<br>with $t^{w2}$ | $p_t$ | No. of pups<br>with $Prl$ indel | $p_c$ |
| --- | --- | --- | --- | --- | --- |
| Male 1 | 30 (4) | 29 | 0.967 | 23 | 0.767 |
| Male 2 | 44 (5) | 42 | 0.955 | 16 | 0.364 |
| Male 3 | 46 (5) | 41 | 0.891 | 41 | 0.891 |
| Male 4 | 46 (6) | 44 | 0.957 | 32 | 0.696 |
| Male 5 | 37 (6) | 37 | 1.0 | 30 | 0.811 |
| **Total** | **203 (26)** | **193** | **0.951** | **142** | **0.7** |
| Female 1 | 12 (2) | 9 | 0.750 | 0 | 0.0 |
| Female 2 | 15 (2) | 5 | 0.333 | 0 | 0.0 |
| **Total** | **27 (4)** | **14** | **0.519** | **0** | **0.0** |

**Table 2:** *Prl* indel frequency, *p*_C_, and predicted loss of function, *p*_L_ from sperm NGS analysis.

| **Mouse** | **$p_C$** | **$p_L$** | **Reads** |
| --- | --- | --- | --- |
| Male 1 | 0.775 | 1.0 | 5045 |
| Male 2 | 0.422 | 1.0 | 4997 |
| Male 3 | 0.819 | 1.0 | 6513 |
| Male 4 | 0.840 | 1.0 | 4767 |
| Male 5 | 0.845 | 1.0 | 6313 |
| Male 6 | 0.874 | 1.0 | 4945 |
| Male 7 | 0.814 | 1.0 | 3967 |
| Male 8 | 0.819 | 1.0 | 3967 |
| Male 9 | 0.882 | 1.0 | 7668 |
| Male 10 | 0.926 | 1.0 | 4245 |
| **Average** | **0.801** | **1.0** | **5243** |

## Discussion

Synthetic gene drives have long been proposed for suppression of invasive alien species and disease vectors. While significant progress has been made in genetic biocontrol of insects, tools are lacking for mammalian pests. Here we describe *t*_CRISPR_, a novel population suppression and even eradication strategy for wild mice. Crucially, our transgenic mouse model demonstrates haplotype transmission, and associated generation and transmission of inactivating mutations in a recessive female fertility gene, all at levels for which our *in silico* modelling predicts that population eradication can occur.

Our model predictions indicate that polyandry has a major impact on spread of *t*_CRISPR_ (and *t^w2^*) and that high levels inhibit *t*_CRISPR_ population suppression. A similar effect has been observed in a recent study of a wild mouse population containing a different *t* haplotype variant^31^. Notably, estimates of multiple paternity in wild mice populations range from 6% - 47%^31,43–47^, which provide reliable lower limits for polyandry estimates; whereas the upper polyandry limit is estimated to be 61% (95% CI: [56% - 65%]) in populations with lethal *t* haplotype^31^. These estimates are within the parameter ranges at which modelling predicts eradication with *t*_CRISPR_, suggesting polyandry may not pose a barrier to achieving this management outcome across a range of realistic scenarios. Additional population genetic data, particularly from island cohorts, would further inform *t*_CRISPR_ eradication potential for wild mice.

Similar to insect homing drives, *t*_CRISPR_ has little tolerance for functional resistance alleles (so called r1 resistance alleles). r1 alleles can likely be limited by using gRNAs that target critical amino acid residues, as we have performed. Alternatively, for fertility genes lacking identifiable critical amino acids, multiple gRNAs could be used. Base editing^48,49^ or prime editing^50^ platforms could also be employed as these can generate desired (loss of function) edits without indels. Interestingly, our modelling indicates *t*_CRISPR_ can cause eradication despite incomplete (e.g. 0.80) cleavage efficiency of the female fertility target gene. Unexpectedly, the *t*_CRISPR_ drive was predicted to be more effective when *p*_C_ = 0.8 than 1.0. Close to 0.80 cleavage was detected in the *t*_CRISPR_ split drive mouse model indicating this is feasible. If required, higher cleavage rates could likely be achieved using other germline active promoters, noting that Cas9 expression does not need to be restricted to the germline as the target fertility gene is not required in males. The *t*_CRISPR_ system is also amenable to multiplexed gRNA approaches targeting multiple fertility genes which may offer protection against resistance allele inhibition.

Restricting gene drive activity to the targeted population is critically important for ensuring no unintended impacts on non-target populations of the same species, for example in their native range^51^. Although islands can provide a natural geographical barrier to unintentional spread, it is generally accepted that multiple safeguards assuring spatial or temporal limitation are required^52,53^. We have recently shown that polymorphisms in female fertility genes that are amenable to SpCas9 cleavage can become fixed in island cohorts^34^. These “locally fixed alleles” could provide potential targets for island-specific *t*_CRISPR_ strategies^53^. Alternatively, integration of the CRISPR cassette in the region immediately flanking the *t* haplotype would allow gradual decoupling from the drive element via meiotic recombination, which may provide a generic strategy for a generation-limited *t*_CRISPR_ drive. Interestingly, our modelling indicates that deployment of *t*^w2^ during *t*_CRISPR_-mediated suppression results in gradual loss of *t*_CRISPR_ from the population suggesting that *t*^w2^ could potentially be used as a natural (non-GMO) reversal drive. Our data also highlights the inhibitory impact of endogenous *t*^w2^ on *t*_CRISPR_-mediated population suppression. Although a *t* variant has been identified in an island population^32^, our bioinformatic screening of island/mainland cohorts suggest that it is not common, consistent with published genetic studies of wild populations.

Our model predictions for the naturally occurring *t*^w2^ are in agreement with earlier models showing that *t*^w2^ can be fixed locally and cause population crashes^25,54^. Compared to *t*_CRISPR_, *t*^w2^ achieved eradication within a much narrower range of parameters and failed as soon as polyandry levels were moderate. Similar to lethal *t* haplotypes^28,29,31^, the frequencies of *t*^w2^ were also reduced with increasing levels of polyandry and sperm competition (Extended Data Fig. 12), which were proposed to be the main contributors to the reduced *t* frequencies observed in nature^31^. Alternative explanations, i.e. unreasonably high fitness costs^55^ and mate choice against *t* haplotypes^56–58^ seem unlikely^31,59^. Encouragingly, *t*_CRISPR_ is effective under a much wider range of model parameters tolerating higher levels of polyandry, sperm competition, and mate choice compared to *t*^w2^.

In summary, we provide *in silico* modelling and transgenic mouse data supporting a unique strategy for suppression and even eradication of invasive mice. Although promising, the impact of many empirical factors remains to be explored in detail including propensity for *t*_CRISPR_ introgression^59–62^, complexity of deme/population structures, seasonal population variation, unanticipated fitness costs and possible DNA target site cleavage resistance due to humoral immunity against Cas9^62^. Although genetic biocontrol strategies including gene drives have considerable potential, their development must proceed with the utmost caution, informed by comprehensive risk assessments, respectful engagement with stakeholders, publics, and the general community, as well as with regulatory authorities^51,63^. We hope that the advances described herein will promote meaningful discussion and debate on the feasibility, benefits and risks of using gene drives to mitigate the devastating impact of invasive rodents on global biodiversity and the environment.

## METHODS

### Animal Models

All experiments involving animal use were approved by the South Australian Health & Medical Research Institute (SAHMRI) Animal Ethics Committee.

### *t*^P2.3-gRNA^ mouse generation

To facilitate integration into the *t* haplotype a *t*^int^-gRNA was designed targeting an intergenic locus adjacent to expressed genes within inversion 1 of the *t* haplotype^41^. Complementary gRNA oligos were designed with overhangs facilitating golden gate assembly and cloned into pSpcas9(BB)-2A-Puro (PX459) V2.0 (Addgene plasmid #62988) according to Ran *et al*. (2013)^64^. Constructs were validated with a Bbsl digest and confirmed by Sanger Sequencing. The gRNA was amplified with oligos containing a 5’ T7 promoter sequence and purified using the QIAquick PCR purification Kit (Qiagen). RNA was generated using the Hiscribe™ T7 Quick High Yield RNA Synthesis Kit (NEB) followed by purification using the RNeasy MinElute Cleanup Kit (Qiagen).

The donor construct was designed using NEBuilder Assembly Tool (https://nebuilder.neb.com/#!/) to facilitate HiFi assembly. pBluescript KS(+) plasmid was linearised with EcoRV and the following four fragments were amplified with HiFi compatible overhangs, i) CMV-mcherry from pZ148-BhCas12b-sgRNA-scaffold (Addgene plasmid #122448), ii) U6-P2.3-HuU6, generated via gBLOCK synthesis (IDT), iii) and iv) *t* haplotype-specific left and right homology arms from *t*/+ mouse gDNA. Each of these fragments were extracted using a DNA Gel Extraction Kit (NEB), quantified, and the HiFi reaction performed as per the manufacturer’s protocol using the NEBuilder HiFi DNA Assembly Cloning Kit (NEB). Constructs were verified by a Sacll and Clal digest followed by Sanger sequencing.

Male *t*/+ mice were mated to superovulated C57BL/6 females and zygotes were harvested. 750ng Cas9 protein (PNA Bio) and 375ng integration site gRNA were mixed, incubated on ice for 10 minutes. 150ng of donor plasmid was added and the total volume made up to 15uL with nuclease-free MQ H_2_O (Invitrogen). This mix was microinjected into the pronucleus of zygotes followed by oviduct transfer into pseudopregnant females. Transgenic founders were identified by mCherry fluorescence and validated by Sanger sequencing of PCR amplicons from using primer pairs flanking the homology arms (Extended Data Fig. 13).

### Tg^Cas9^ mouse generation

The SpCas9-EGFP-bGH poly(A) cassette from Addgene plasmid #61408 was cloned into the pStart-K vector (Addgene #20346). A *Ccna1* promoter region extending 5.7 kb upstream of the start codon (exon 2) was amplified by PCR from C57BL/6 mouse genomic DNA. This region was cloned 5’ of Cas9 in the pStart-K-Cas9-EGFP backbone. An 11.8 kb fragment containing the *Ccna1* promoter fragment, Cas9, EGFP and bGH poly(A) sequence was digested, gel extracted and purified. Forty five nanograms of this fragment in 15 uL of 1x microinjection buffer was injected into the pronucleus of zygotes and transferred into pseudopregnant females. A single transgenic founder was confirmed by PCR, and further characterised by immunofluorescence, qPCR and digital PCR (Extended Data Fig. 11).

### Immunofluorscence

Testis immunofluorescence was performed as described^65^ using a chicken anti-GFP primary antibody (Abcam, #ab13970) diluted 1/600 and an anti-chicken Alexa Fluor 488 secondary antibody (Jackson ImmunoResearch #103-545-155) diluted 1/300.

### *t*_CRISPR_ split drive analysis

Double hemizygous *t*_CRISPR_ split drive males were generated by crossing *t*^PZ3-gRNA^ and Tg^Cas9^ mice. *t*_CRISPR_ males were housed with two wild type C57BL/6 females, who were checked daily for vaginal plugs. Plugged females were removed and replaced with a new wild type female. Embryos were harvested from pregnant females at 13.5-17.5 dpc. Embryonic DNA was extracted using the KAPA Express Extraction kit (Roche) according to the manufacturer’s instructions. The P2.3 gRNA target site was amplified, and PCR products purified by the QIAquick PCR Purification Kit (QIAGEN) and Sanger sequenced. Indel analysis was performed using DECODR (https://decodr.org/). *t* haplotype transmission was assessed by genotyping the Hba-ps4 locus of the *t* allele as described by Schimenti and Hammer (1990)^66^. Female *t*_CRISPR_ mice were mated with C57BL/6 males. Ear biopsies from pups were taken at P10 and indel and *t* haplotype transmission was analysed as above.

To assess *Prl* indel generation in the *t*_CRISPR_ male germline, sperm from *t*^P2.3-gRNA/0^/Tg^Cas9/0^ mice was isolated from the cauda epididymis using the swim up method. Briefly, extracted sperm was pelleted by centrifugation for 10 minutes at 360g. The supernatant was removed and replaced with 10 FCS + DMEM followed and the sperm repelleted. The tube containing the sperm was placed at a 45° angle in a 37°C, 5% CO_2_ incubator for 1h followed by removal of the sperm-containing supernatant and centrifugation. Sperm DNA was extracted using the High Pure PCR Template Preparation Kit (Roche) modifying the first step of the protocol. Briefly, pelleted sperm were resuspended in 400uL tissue lysis buffer and 50uL of proteinase K and incubated for 1h at 55°C. 50uL of 1M DTT was added, mixed, and incubated overnight at 55°C. 40uL of proteinase K was added and the protocol was continued as per manufacturer’s instructions, eluting in 100uL of elution buffer.

### Cell culture and Transfection

Complementary gRNA oligos were designed with overhangs facilitating golden gate assembly and cloned into pSpCas9(BB)-2A-Puro (PX459) V2.0 (Addgene plasmid #62988) according to Ran *et al*. (2013)^64^. R1 Mouse embryonic stem cells were cultured and transfected as described by Adikusuma *et al*. (2021)^67^ without modification.

### Next Generation Sequencing

DNA extracted from *t*^P2.3-gRNA/0^/Tg^Cas9/0^ sperm or mES cells was amplified with Nextera-tagged primers under standard Phusion PCR conditions. PCR products were sent to the Australian Genome Research Facility for barcoding and paired end NGS using the MiSeq Nano (500 cycles). NGS reads were analysed using CRISPResso2^68^ using default parameters except: Quantification window size (5bp), minimum average read quality >20, minimum single bp quality >10 and replace bases with N that have a quality lower than <10.

### DNA sequences

Oligonucleotide sequences are listed in Extended Data Table 2.

### *In silico* modelling

The individual-based and spatially explicit modelling framework presented in Birand *et al*. (2022)^69^ was used. This is a stochastic, discrete-time model with overlapping generations. Individuals are diploid, have genetically controlled autosomal traits and sex chromosomes, and occupy a rectangular array of patches that together form a hypothetical island. Patches hold multiple individuals and individuals can use multiple patches. Each breeding cycle is considered a model time step and individuals that survive long enough pass through a number of breeding cycles until they reach a maximum age (*age*_m_). The number of breeding cycles per year is given by *n*_c_, with the following steps occurring each cycle (for more details, see Birand *et al*. (2022)^69^):

1. Mate search: An individual’s mate search area is determined by all the patches surrounding its central patch within distance *D*_m_. The search for mates begins within the central patch and continues incrementally with distance in the neighboring patches until a mate is found. All patches of equal distance have the same probability of being chosen during the mate search. Individuals retain their central patch within a breeding cycle, irrespective of whether they find a mate.
2. Mating. All females mate unless there are no males present in patches within distance *D*m, or if they are infertile due to the lack of a functional fertility gene. Since females choose males randomly, some males can mate multiple times in a single breeding cycle, whereas others may not mate at all. Polyandry is permitted within a single breeding cycle; when multiple males are present in a female’s central patch, the female can mate with multiple males (*n*_m_) according to the probability of multiple mating (*p*_m_)^31,44^. To account for the possibility that females with the *t* haplotype might prefer to mate with wild type males, the probability of these females selecting a *t*-carrying male is reduced by an aversion coefficient (*a*_c_).
3. Density-dependent reproduction. The number of offspring from each mated female is drawn from a Poisson distribution with mean *u* = *b*/(1 + [(*b*/2) - 1][*N*/*K*]) (discrete-time Beverton-Holt model^70^), where *b* is the average number of offspring of females in the absence of density-dependent regulation, *N* and *K* are the population size and the carrying capacity in a female’s central patch, respectively. Under polyandrous mating, the paternity of each offspring is determined randomly based on a probability assigned to each of the possible fathers. For *t* haplotype carrying males, the probability of siring an offspring is reduced compared to that of the wild type male by a sperm-disadvantage coefficient *d*_s_. Studies show the majority of litters showing multiple paternity are sired by two males (~80-85%)^31,45,46^. If the female mates with two males, the coefficient range between 0 and 0.5; *d*_s_ = 0 means that both males have equal probability of siring the offspring, whereas *d*_s_ = 0.5 means that wild type male will sire the offspring (corresponds to competitive skew = [0.5, 1] in Dean *et al*. (2006)^44^, and relative competitiveness, *r* = [1,0] in Manser *et al*. (2017, 2020)^29,31^). *d*_s_ = 0.17 mean that probabilities of siring an offspring 0.33 (0.5 - 0.17) and 0.67 (0.5 + 0.17), for the *t* carrying male and wild type male, respectively, which could be the case if the former has half the sperm amount compared to the latter (*r* = 0.5 in Manser *et al*. (2020)^31^). Values *d*_s_ > 0.17 correspond to greater sperm disadvantages that cannot be explained by reduction of sperm amount alone. The sex of each offspring is determined by the sex chromosomes inherited from parents.
4. Natal dispersal. All offspring are assumed to survive and become subadults since offspring mortality is incorporated in the density-dependent reproduction function above. A subadult can disperse to a new patch from its maternal patch within distance *D*_n_ to establish a mate-search area before its first breeding attempt. Natal dispersal is both distance and negative density dependent^69^, which ensures that when the population size is close to carrying capacity, the probability of dispersing long distances is low. Modifying this assumption to model random or positive density dependent dispersal does not substantially impact the simulated probability of eradication or time to eradication^69^.
5. Survival of adults. Adult survival probability (*ω*) is constant for each breeding cycle. To incorporate the possibility of an additional fitness cost, the survival probability of *t* haplotype carrying males is reduced further by a multiplicative constant *ω*_t_.
6. Breeding dispersal. Surviving adults can establish new mate-search areas with a new central patch within distance *D*_b_^71^. The probability of moving a distance *δ* is calculated as natal dispersal above (for simplicity, we assume that the maximum distances for natal dispersal, breeding dispersal, and mate-search distance parameters are physiologically constrained to be equal, and determined by *D*). The negative density-dependent dispersal function ensures that when the population size is close to carrying capacity, individuals (subadults or adults) tend to retain the same central patch for natal and breeding dispersal, and form long-term stable communities^44,72,73^. At low densities, the probability of picking a distant patch to centre its new mate-search area is higher, which can be justified by an individual’s imperative to move large distances to find mates when local density is nearly zero^69,74–76^.

#### Suppression and eradication strategies

The *t* haplotype is a naturally occurring segregation distorter with high transmission rates (*p*_t_) to offspring in heterozygous males. The *t*^w2^ variant causes sterility in homozygous males but has no inheritance-biasing or fertility effect in females. We explored the effects of a Cas9/gRNA construct embedded in this variant (hereafter *t*_CRISPR_) targeting a fertility gene (e.g. prolactin *Prl*) located on another autosomal chromosome. The fertility gene is present in both sexes but required only in females. The gene is haplosufficient, meaning that when both copies of the gene are deactivated, the female is infertile. We initially assumed that Cas9/gRNA is active in the germline of males only, but also performed simulations when it is active in the germline of both sexes (denoted *t*_CRISPR(2)_).

We explored the possibility of evolution of resistance when the fertility gene is cleaved by Cas9/ gRNA. The probability of successful fertility gene knockout is given by *p*_C_*p*_L_, where *p*_C_ is the probability of a successful DNA cut by Cas9/gRNA, and *p*_L_ is the probability of loss of gene function following a successful cut. We assumed that if the fertility gene retains its function after a cut, it is also resistant to further cuts; therefore, the probability that the resulting allele is functional and resistant is *p*_C_(1 – *p*_L_). The probability of a failed knockout is (1 – *p*_C_), and the allele remains functional and susceptible to further cuts.

We also explored the effect of releasing only naturally occurring *t*^w2^ variant males without a Cas9/gRNA construct and checked its effectiveness as a population suppression tool via male sterility.

Lastly, we compared the effectiveness of *t*_CRISPR_ as a suppression and eradication strategy with the homing and X-shredding drives, which were investigated in detail by Birand *et al*. (2022)^69^. The homing drive is a CRISPR-based drive that is positioned in an exon of the same female fertility gene targeted by *t*_CRISPR_ and generates a loss-of-function mutation. The X-shredding drive is a CRISPR-based drive located on the Y chromosome and cuts the X chromosome with probability *p*_x_ at multiple locations beyond repair during spermatogenesis. Since X-bearing sperm are destroyed, and eggs are predominantly fertilized by Y-bearing sperm, the drive causes disproportionately more male offspring.

#### Parameters and initial conditions

Life history and demography parameters (Extended Data Table 1) are based on empirical data whenever available (for details, see Birand *et al*. (2022)^69^). We use the same spatial configuration provided in Birand *et al*. (2022)^69^: 64 x 64 = 4096 patches in the model corresponds to a 70m x 70m space on a hypothetical island of approximately 2000 ha. We calibrated dispersal abilities on this island based on historical invasion records of a similar sized island^77^; maximum dispersal distances *D* = [4,7] in the model correspond to 240 – 560m in the wild^74,78^. We assumed that the population size was ~200,000 before inoculation.

After a burn-in period of two years, the simulated island is inoculated with males carrying *t* haplotype variants (*t*_CRISPR_ or *t^w2^*) detailed above. We modelled a single release into 256 patches distributed systematically across the island, with one male (*N*_i_ = 1) with *t*_CRISPR_ or *t*^w2^ released into each patch (total of 256 males), assuming that would be an achievable release size. Releasing more individuals with gene drives reduces the simulated time to eradication (e.g. Birand *et al*. (2022b)^79^). Following their release, *t*-carrying mice randomly choose a patch within distance *D* from their inoculation patches and join the pool of available males for mating. Females choose males randomly among all the available males, which implies that some *t* haplotype carrying males may not be chosen for mating (potentially more so if there is mate choice against *t* haplotype males by *t* haplotype females).

We ran simulations for a maximum of 500 breeding cycles (i.e., 81 years after inoculation) and compared the efficacy of different gene-drive strategies under different parameter assumptions. We also performed a global sensitivity analysis for each suppression strategy to investigate the relative influence of parameters on the probability of successful eradication and the time to eradication (see below), in addition to brute force simulations with certain parameter combinations. The model is coded using C programming language and is available from the Zenodo Open Repository (10.5281/zenodo.6583793).

#### Sensitivity analysis

We performed a global sensitivity analysis for each suppression strategy in order to investigate the relative influence of parameters on the probability of successful eradication and the time to eradication. In each case, we created 3,000 unique parameter combinations from parameter ranges given in Extended Data Table 1 using Latin hypercube sampling (randomLHS, R package^80^). In order to minimize computational effort and maximize the coverage of the parameter space, we ran a single simulation for each parameter combination^81^. Finally, we examined the influence of parameter inputs using Boosted Regression Tree models (BRT; R package *dismo*^82^) that we fitted to the simulation outputs using the function gbm.step from the R package *dismo* (learning rate: 0.01; bag fraction: 0.75; tree complexity: 3; and 5-fold cross-validation^83^).

We used binomial error distribution for the probability of eradication, and Poisson error distribution for time to eradication^83^. We labelled simulation outcomes as unsuccessful for the binomial error distribution if eradication did not occur within the number of breeding cycles simulated, even when the population was suppressed to a new stable level. To investigate the influence of parameters on the time to eradication, we only used simulations when eradication was achieved.

### Presence of *t* haplotypes on islands

Given the potential to undermine *t*_CRISPR_ efficacy, knowledge of *t* haplotype prevalence in island populations is critical, yet information from previous studies has been sparse, contradictory, and often based on small sample sizes. For example, early surveys of North American house mouse populations noted several island populations among the rare exceptions where *t* haplotypes were not detected^26,84,85^. A later study of isolated island populations in Great Britain identified a *t* haplotype present on one island, but absent in two others^32^. More recently, whole-genome sequencing of *M. m. helgolandicus* from the island of Heligoland found no evidence of *t* haplotypes, though only a small number of individuals were sampled (N=3)^86^.

To assess the presence of *t* haplotypes in island mice, we analysed previously published pooled whole-genome sequences (pool-seq) from four introduced populations on islands in Western Australia (Whitlock-Boulanger Islands and Thevenard Island), the northern Pacific (Sand Island in Midway Atoll), and North America (South Farallon Island)^34^. As it is generally not possible to extract haplotype information from pool-seq data, we used an approach wherein sequencing reads were mapped to a reference of diagnostic *t*-specific transcripts. These reference transcripts were derived from results of a recent study by Kelemen *et al*. (2022)^87^ that analysed DNA and RNA sequences from a database of wild *Mus* samples^86^. This database includes whole-genome sequences from both +/*t* and +/+ mice across all three *M. musculus* subspecies as well as a sister taxon (*M. spretus*). The original list (Supplementary Data 4 in Kelemen *et al*. (2022)^87^ consists of 256 annotated contigs assembled from RNA-seq reads of +/*t* mice, and subsequently filtered for those that had greater relative coverage of genomic reads mapped from +/*t* compared to +/+ mice.

In order to identify diagnostic *t*-specific transcripts for our study, we downloaded DNA sequence FASTQ files for +/*t* and +/+ mice from the Harr *et al*. (2016)^86^ database and re-mapped the forward reads of each sample to the transcripts using bowtie 2^88^. Following the procedure by Kelemen *et al*. (2022)^87^, we then counted only reads that mapped perfectly to a transcript with no mismatches, and normalised these values by the total number of reads in each sequence dataset, multiplied by 1 x 10^6^ to calculate reads per million (RPM). Diagnostic *t*-specific transcripts were then identified as those to which zero reads mapped from any +/+ individuals, and at least one read mapped in each +/*t* sample. This final list of transcripts was then used to screen island pool-seq data for the presence of *t* haplotypes in each population. At the same time, we also analysed the four ‘source’ mouse populations from the same study as a control^34^, as some have speculated that *t*-alleles may tend to be more common in larger populations that are less influenced by genetic drift^89^. We downloaded the FASTQ files for each pool-seq dataset from the NCBI Short Read Archive (Bioproject PRJNA702596) and mapped the first reads to the set of diagnostic *t*-transcripts as above, and calculated the fraction of transcripts with at least one read mapped as well as the normalized counts of reads mapped.

As our primary aim was to evaluate presence or absence of *t* haplotypes in each island population, we compared observed *t*-specific read counts to the expected values under a scenario where a single *t* haplotype-bearing chromosome was sampled within the pool-seq experiment. To simulate this, we concatenated FASTQ files from +/*t* samples in the Harr *et al*. (2016)^86^ database, and randomly subsampled using rasusa^90^ down to a coverage equal to 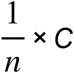, where *n* and *C* are the number of haploid genomes sampled in each empirical pool-seq sample and the total library sequencing coverage (first reads only), respectively (see Table S3, Oh *et al*. (2021)^34^). The process was repeated with a concatenated FASTQ file of reads from +/+ samples, subsampled to 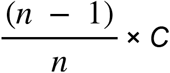, then both files were combined and mapped to the diagnostic *t*-specific transcripts as above to generate expected counts. Each simulation was iterated 100 times in each population to generate mean expected values and 95% confidence intervals.

We note that our analyses assume equal contribution of each individual sample within the pool-seq libraries. Violation of this assumption could lead to under- or overrepresentation of reads mapping to *t*-specific transcripts in the data, though the original study^34^ adopted best practices^91^ to minimise such biases (e.g. DNA was quantified fluorometrically in triplicate prior to pooling).

## Acknowledgements

The findings and conclusions in this publication are those of the authors and should not be construed to represent any official USDA or U.S. Government determination or policy. This research was supported by the Government of New South Wales, the Centre for Invasive Species Solutions and the Government of South Australia. This research was also supported in part by the U.S. Department of Agriculture, Animal Plant Inspection Service, Wildlife Services, National Wildlife Research Center. The authors acknowledge the facilities and the scientific and technical assistance of the South Australian Genome Editing (SAGE) Facility, the University of Adelaide, and the South Australian Health and Medical Research Institute. SAGE is supported by Phenomics Australia. Phenomics Australia is supported by the Australian Government through the National Collaborative Research Infrastructure Strategy (NCRIS) program. This work was supported with supercomputing resources provided by the Phoenix HPC service at the University of Adelaide. We thank Dan Tompkins and members of the GBIRd consortium for commenting on the manuscript.

## Author Contributions

P.Q.T. conceived the study with input from J.G. and D.W.T. A.B. performed the in silico modelling analysis with input from T.A.A.P. L.G. performed the cell culture and mouse experiments with assistance from M.D.B., G.I.G., S.G.P. K.P.O. performed the island cohort bioinformatic analysis with input from A.J.P. P.Q.T, T.A.A.P., P.C., J.V.R. and O.E. supervised the study and all authors contributed to data analyses. P.Q.T., A.B. and L.G wrote the manuscript. All authors participated in editing the manuscript.

## Code availability

The code for the individual-based model is available from the Zenodo Open Repository (10.5281/zenodo.6583793).

## Data availability

Additional data that support the findings of this study are available from the corresponding author upon reasonable request.

Supplementary Information is available for this paper.

## EXTENDED DATA

**Extended Data Table 1.** Parameters of the model. Base values are used as default values in most of the simulations. If other values are used, they are mentioned in the figure captions. For sensitivity analyses (SA), parameter combinations are drawn from a uniform distribution (*U*) or uniform discrete distribution (*U*_d_) using Latin hypercube sampling.

| Parameter | Base value | SA |
| --- | --- | --- |
| *Life history:* |  |  |
| average number of offspring ($b$) | 6 | |
| maximum age ($age_m$) | 2 | |
| number of breeding cycles in a year ($n_c$) | 6 | |
| dispersal and mate-search distance ($D$) | 5 | $U_d(4, 7)$ |
| relative fitness of individuals with $t$ haplotype ($\omega_t$) | 1 | $U(0.9, 1)$ |
| probability of survival ($\omega$) | 0.53 | $U(0.40, 0.66)$ |
| carrying capacity per patch ($K$) | 19 | |
| *Mating:* |  |  |
| probability of multiple mating ($p_m$) | 0.46 | $U(0.2, 0.8)$ |
| number of males mated per breeding cycle ($n_m$) | 2 | |
| sperm-disadvantage coefficient ($d_s$) | 0.17 | $U(0.17, 0.39)$ |
| aversion coefficient ($a_c$) | 0 | |
| *Inoculation:* |  |  |
| number of inoculation sites | 256 |  |
| number of gene-drive carrying individuals inoculated ($N_i$) | 1 | |
| number of releases over time ($n_t$) | 1 | |
| *$t$ haplotype parameters:* |  |  |
| transmission probability of $t$ haplotype in heterozygous males ($p_t$) | 0.95 | $U(0.93, 0.99)$ |
| probability of successful cut ($p_c$) | 0.8 | $U(0.6, 0.9)$ |
| probability of loss of function after a successful cut ($p_L$) | 1.0 | $U(0.94, 0.96)$ |

**Extended Data Table 2.**
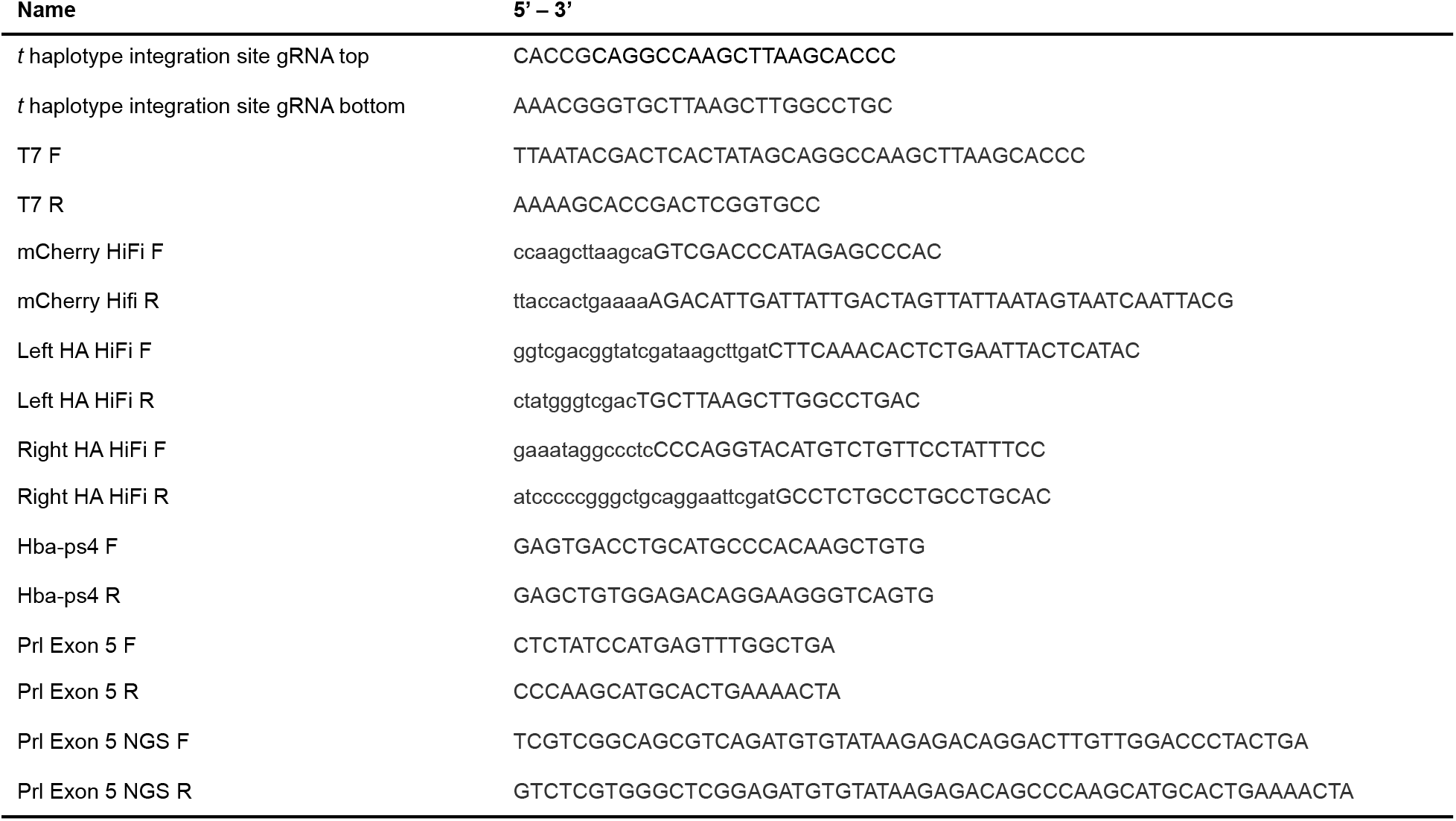
Oligonucleotides used in this study.

**Extended Data Figure 1.**
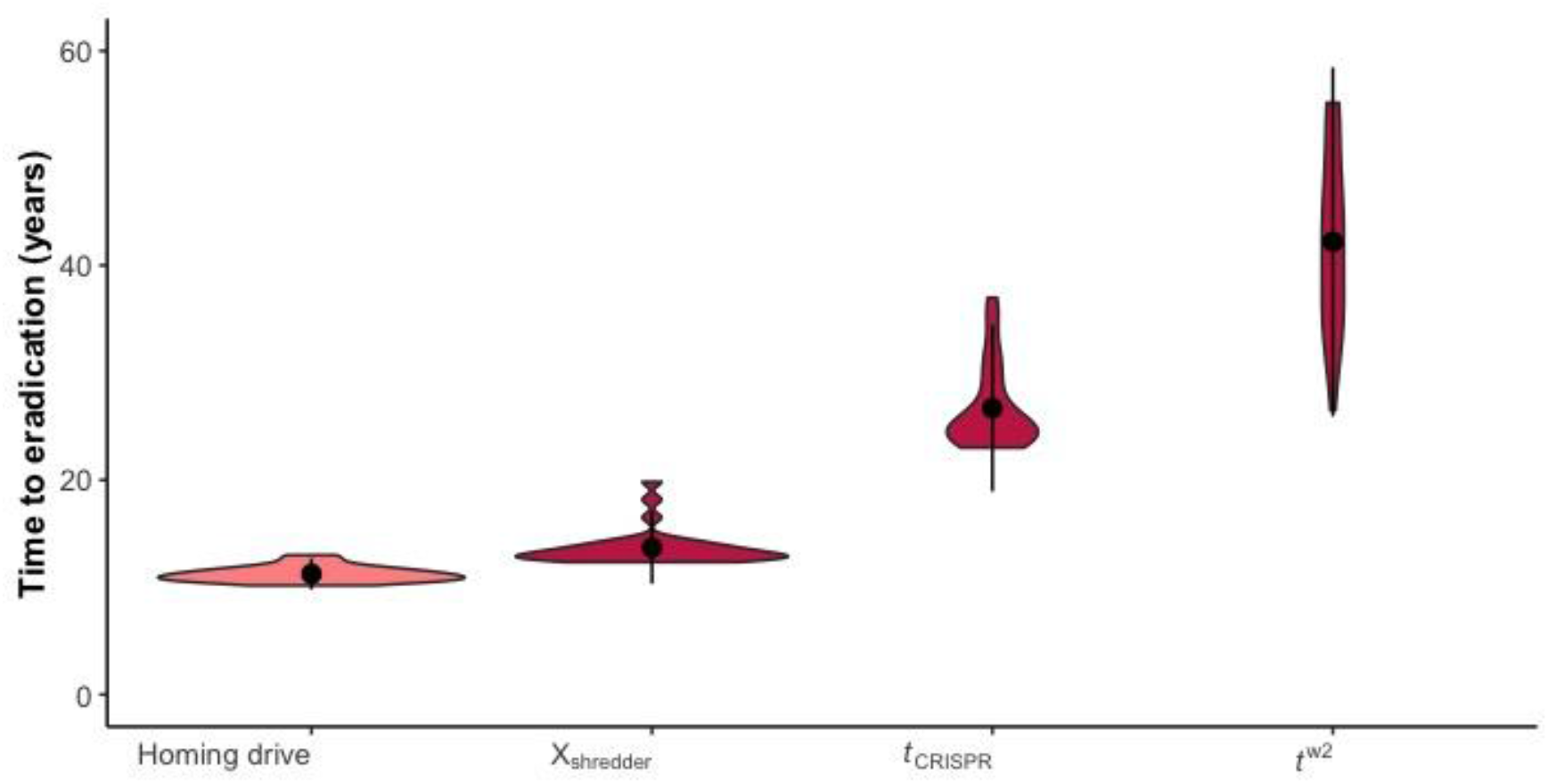
Violin plots showing the density distribution of time to eradication in 30 simulations for each strategy (mean and two standard deviations as black circle and lines). For homing drive, we assumed that the probability of a successful DNA cut, *p*_C_ = 0.90, the probability of Non Homologous End Joining, *p*_N_ = 0.01, and the probability of loss of gene function following NHEJ, *p*_L_ = 0.9999. For the X-shredder, the probability of Y-drive shredding the X chromosome, *p*_x_ = 0.96. Colors represent the probability of eradication, which was 0.7 in the homing drive and 1 in the others.

**Extended Data Figure 2.**
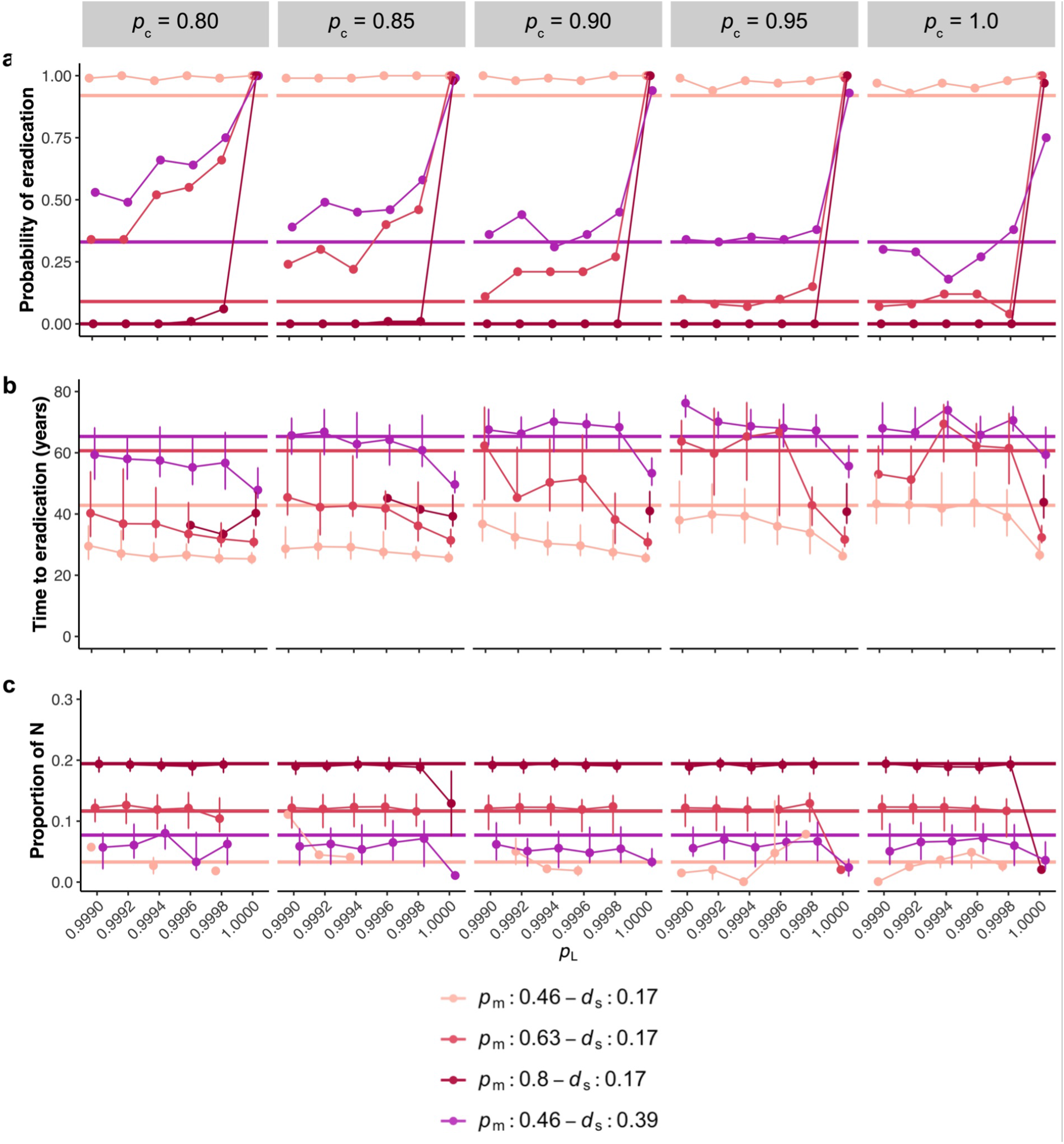
The effect of probability of successful DNA cut, *p*_C_, and loss of function after a cut, *p*_L_, using *t*_CRISPR_ when *p*_t_ = 0.95 compared against *t*^w2^ (solid horizontal lines) with various levels of polyandry and sperm competition. **a**. The probability of eradication; the median with interquartile ranges of the time to eradication (**b**), and the proportion of remaining population sizes when eradication was unsuccessful (**c**) based on 100 simulations for each parameter combination (12000 simulations in total).

**Extended Data Figure 3.**
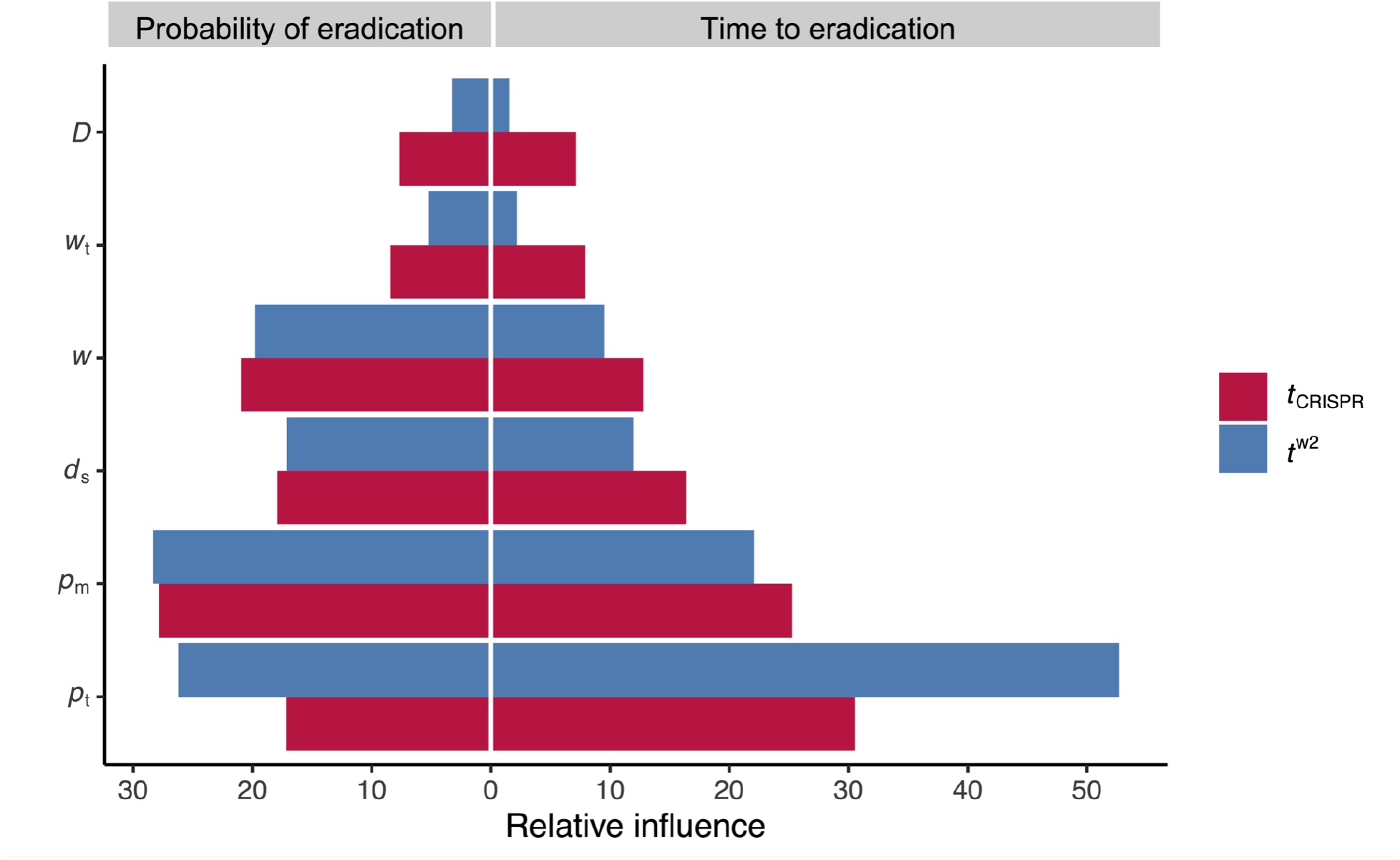
Relative influence of parameters on the probability of eradication (left) and the time to eradication (right) using *t*_CRISPR_ and *t*^w2^ from Boosted Regression Tree models fit to the sensitivity-analysis output (based on 3000 simulations for each strategy for probability of eradication; 1641 and 1634 simulations for *t*_CRISPR_ and *t*^w2^, respectively, for time to eradication where eradication was successful.) Relative influence of *p*_C_ and *p*_L_ were less than 1% each, when included in the sensitivity analysis (based on another set of 3000 simulations, results not shown).

**Extended Data Figure 4.**
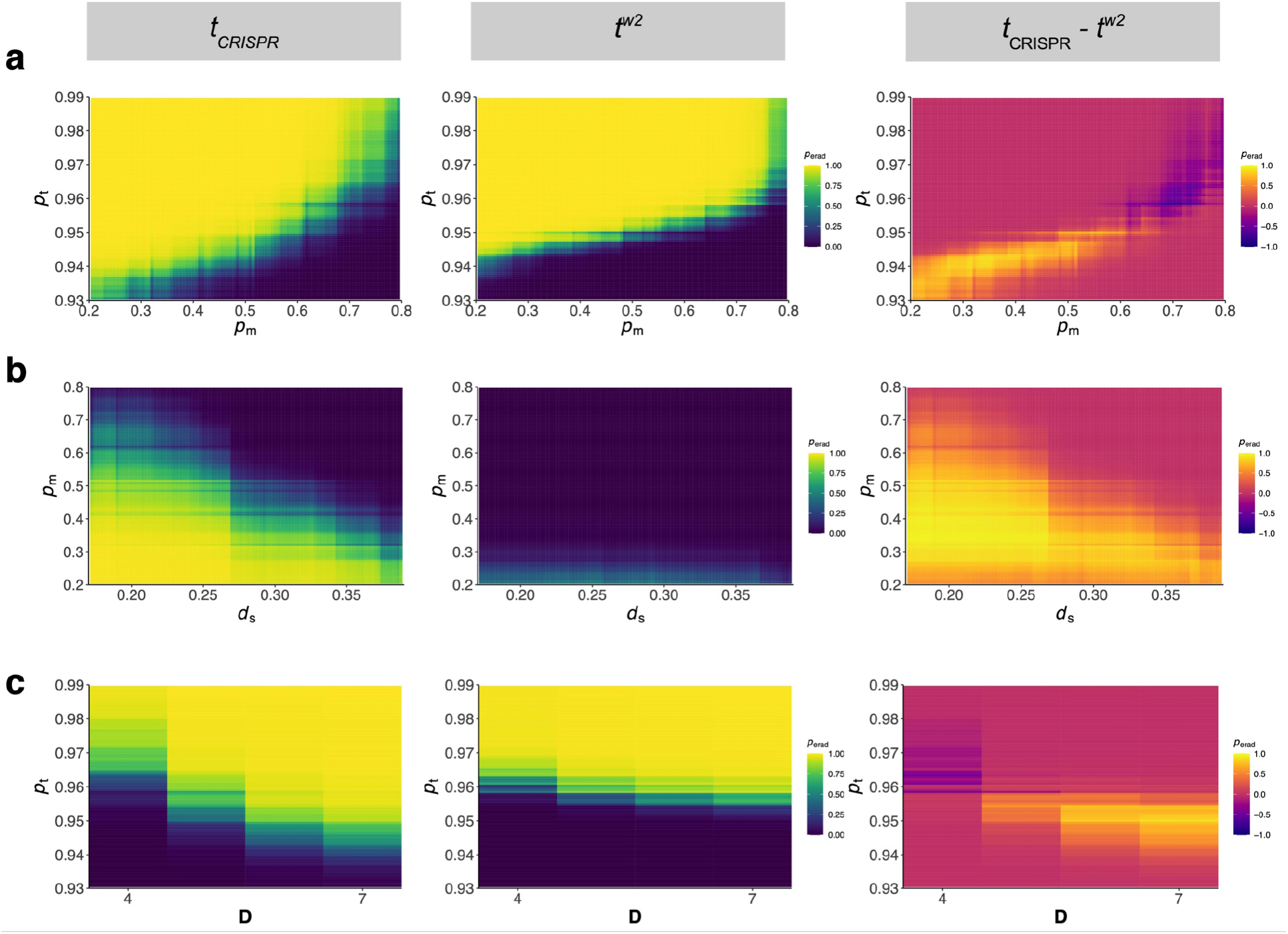
Refer to the caption of Figure 4 in the main text. We assumed that **a**. high sperm competition *d*_s_ = 0.28; **b.** *p*_t_ = 0.94; and **c.** *p*_m_ = 0.8.

**Extended Data Figure 5.**
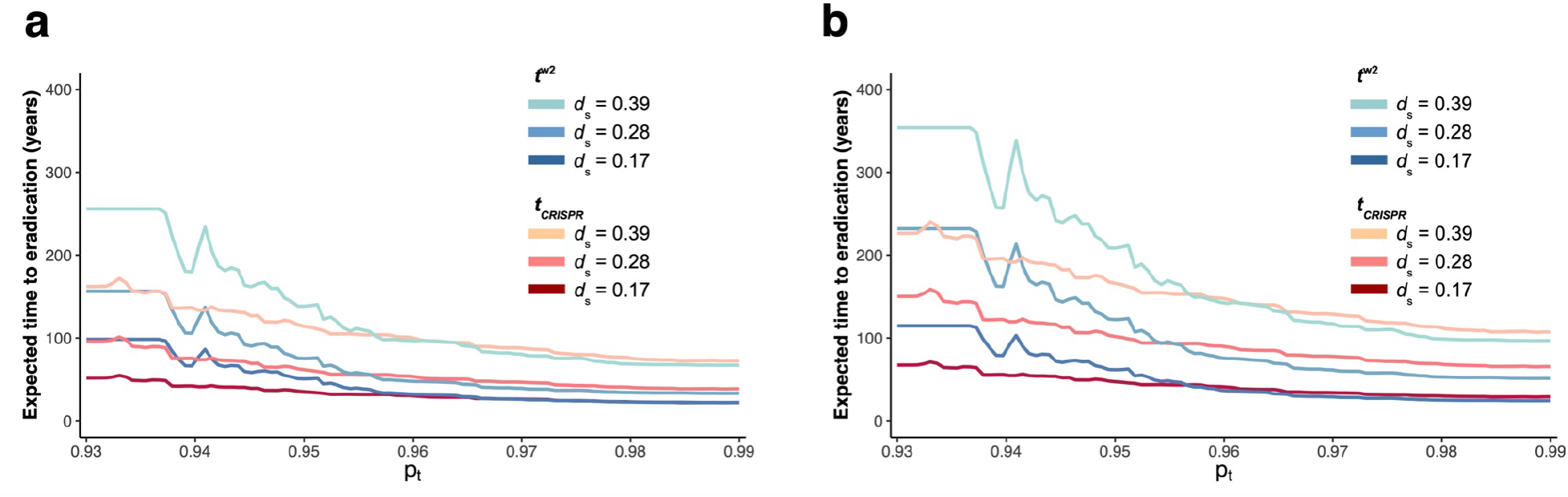
Refer to the caption of Figure 5a in the main text **a**. *p*_m_ = 0.63, **b**. *p*_m_ = 0.8.

**Extended Data Figure 6.**
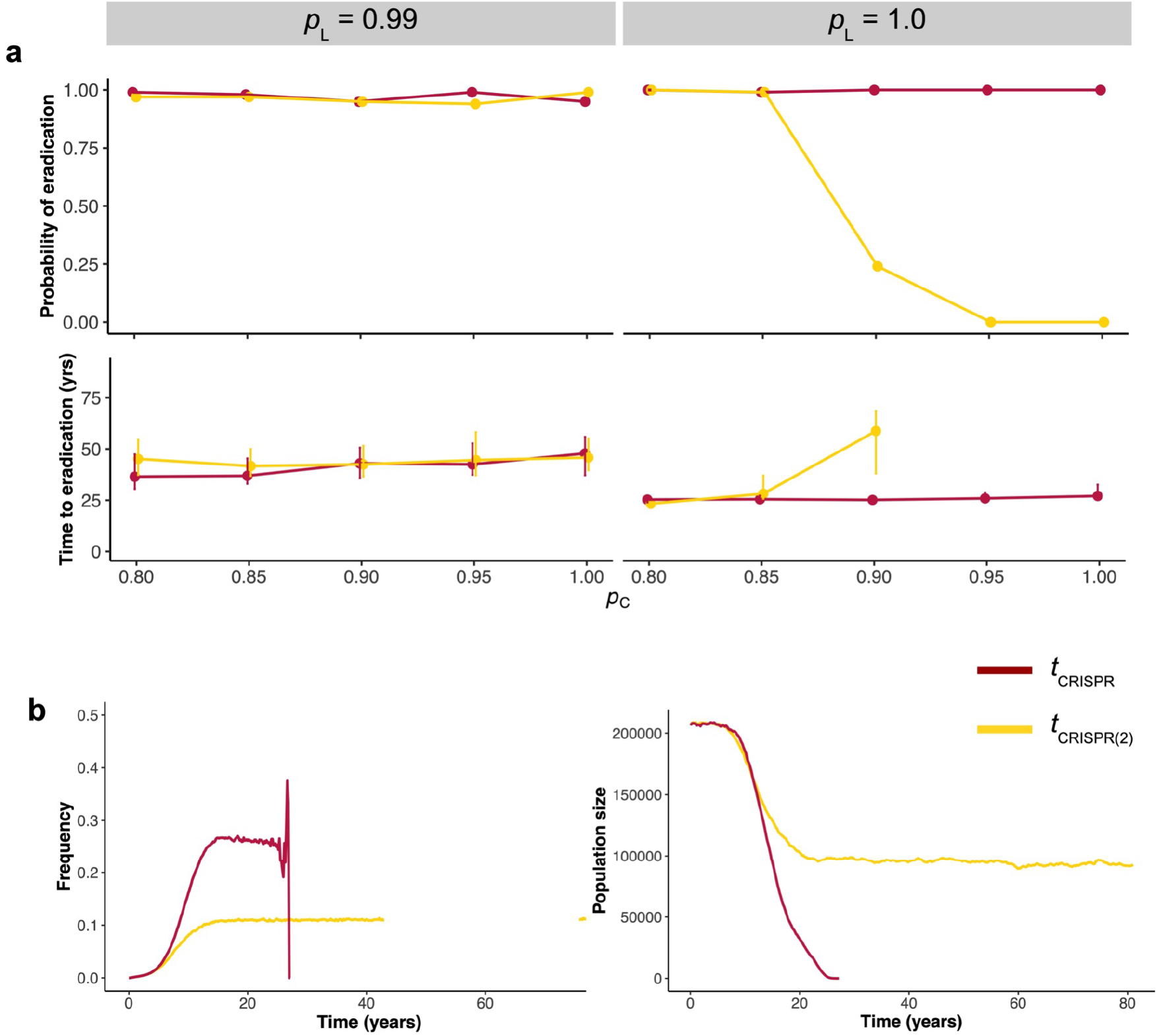
**a**. Probability of eradication and time to eradication using *t*_CRISPR_, where *Prl* knockout is in male germline only; and with *t*_CRISPR(2)_, where *Prl* knockout is in the germline in both sexes with various probabilities of successful cut *p*_C_ and loss of function after a cut *p*_L_ (based on 100 simulations for each parameter combination, 2000 simulations in total, assuming *p*_m_ = 0.46). **b**. Frequencies of fertile females with the *t* haplotype (left) and population size through time (right) in two sample simulations with *p*_C_ = 1 and *p*_L_ = 1.

**Extended Data Figure 7.**
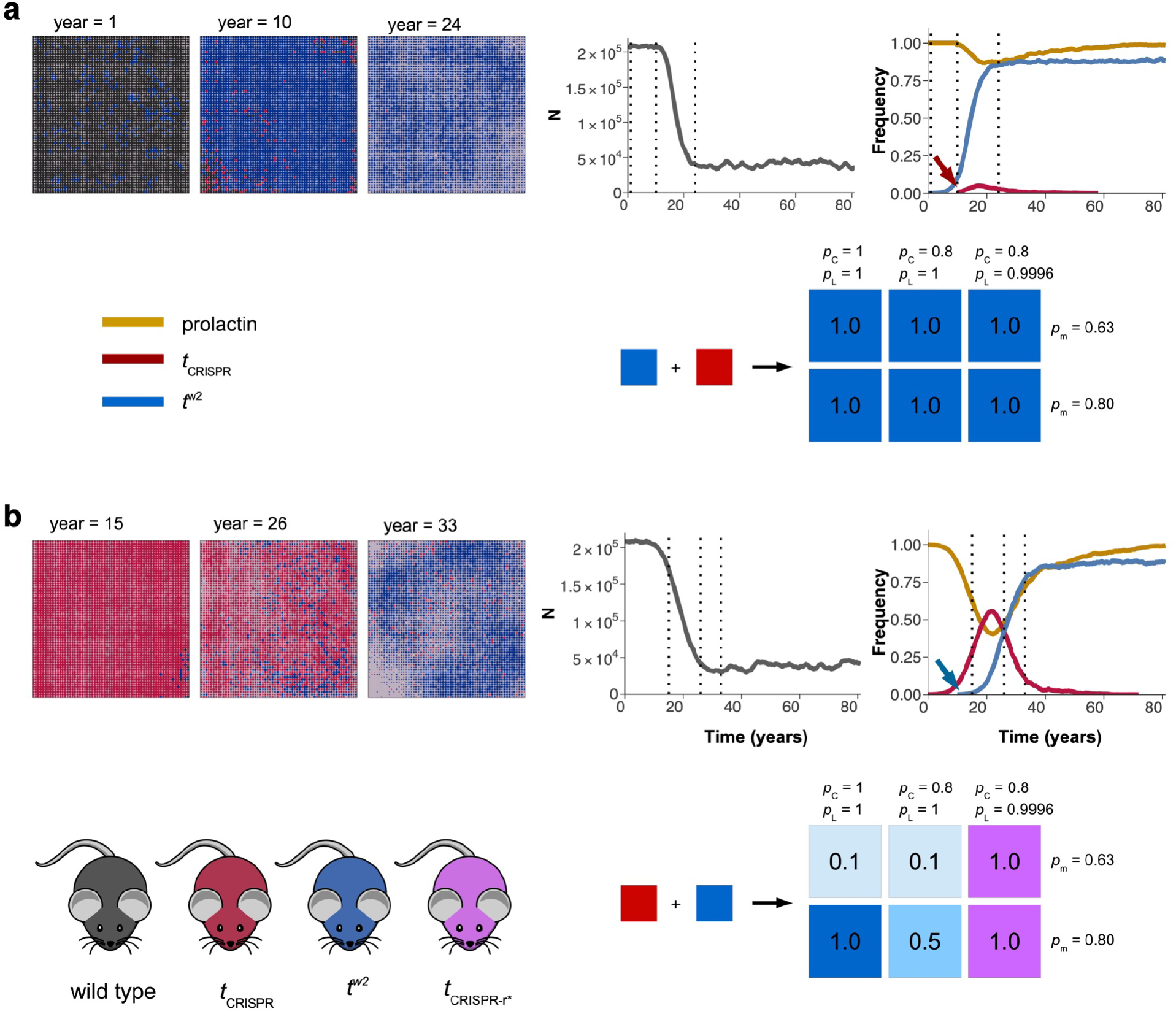
Dynamics of multiple *t* haplotypes (refer to the caption of Figure 2 in the main text). **a.** A sample simulation showing the failure of *t*_CRISPR_ (inoculation at year 10, red arrow) to invade when the island was already inoculated with *t*^w2^ individuals at year 0. Even though the *t*_CRISPR_ inoculation was done with 10 individuals in each of the 256 inoculation patches, it failed to spread, and disappeared from the population in ~50 years after inoculation. The result was the same under various parameter combinations where *t*_CRISPR_ disappeared and *t*^w2^ remained in all simulations (lower plot with probabilities). **b.** A sample simulation showing potential “rescue effect” by *t*w^2^ (inoculation at year 10, blue arrow line), after the island was inoculated with *t*_CRISPR_ individuals at year 0. Five years after *t*^w2^ inoculation with 1 individual/patch at year 10, *t*_CRISPR_ was still the most abundant haplotype in the population, but it disappeared from the population in < 80 years. Rescue effect was sensitive to parameters, and had a higher probability of being successful when polyandry was high (lower plot). For most of the values, rescue failed either because the population went extinct, or resistant *t*_CRISPR-r_* evolved. (Both sample simulations with *p*_m_ = 0.8, *p*_C_ = 1, and *p*_L_ = 1; based on 10 simulations for each parameter combination).

**Extended Data Figure 8.**
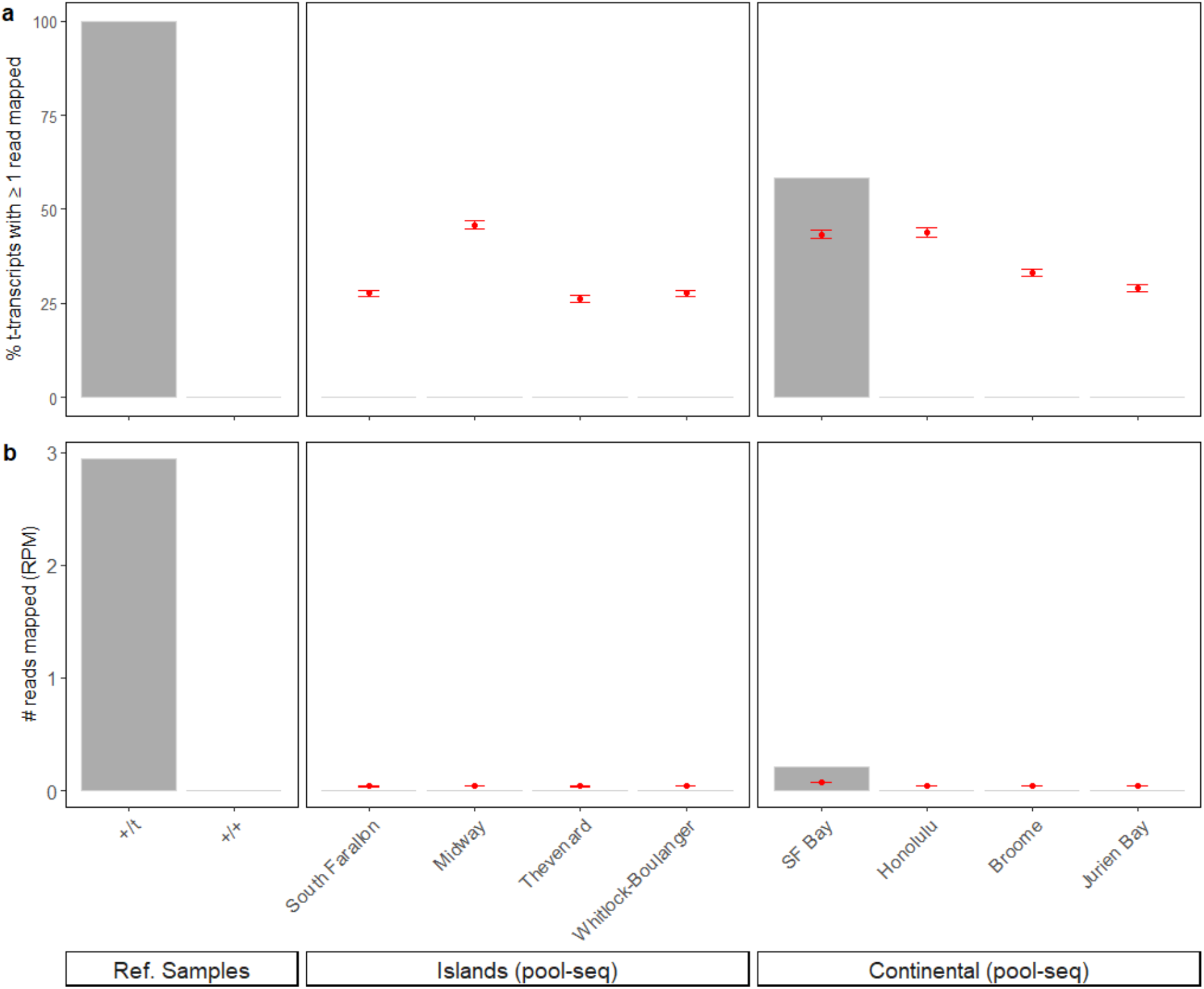
Mapping of genomic sequence reads to 55 diagnostic transcripts (adapted from Kelemen *et al*. (2022)^87^) to detect *t* haplotypes in mouse pool-seq datasets (Oh *et al*. (2021)^34^). Diagnostic *t*-specific transcripts were identified by first mapping reads from *t*/+ and +/+ reference samples (left panel)^86^ and calculating (a) the percentage of diagnostic transcripts to which one or more genomic reads mapped perfectly, and (b) the total number of reads mapping perfectly to diagnostic transcripts, normalised to total dataset size and scaled to reads per million (RPM). Analysis of pool-seq data from small, isolated island populations (center panel) and three of four ‘continental’ populations (right panel) showed no evidence of *t* haplotypes; whereas one population (SF Bay) had genomic reads consistent with at least one *t* haplotype within the pooled sample. Red points are the mean expected values and 95% confidence intervals for each population at minimal detection limits (i.e., sampling of a single *t*/+ individual) in simulated datasets (100 iterations) with equivalent parameters to the original empirical dataset (refer to Methods).

**Extended Data Figure 9.**
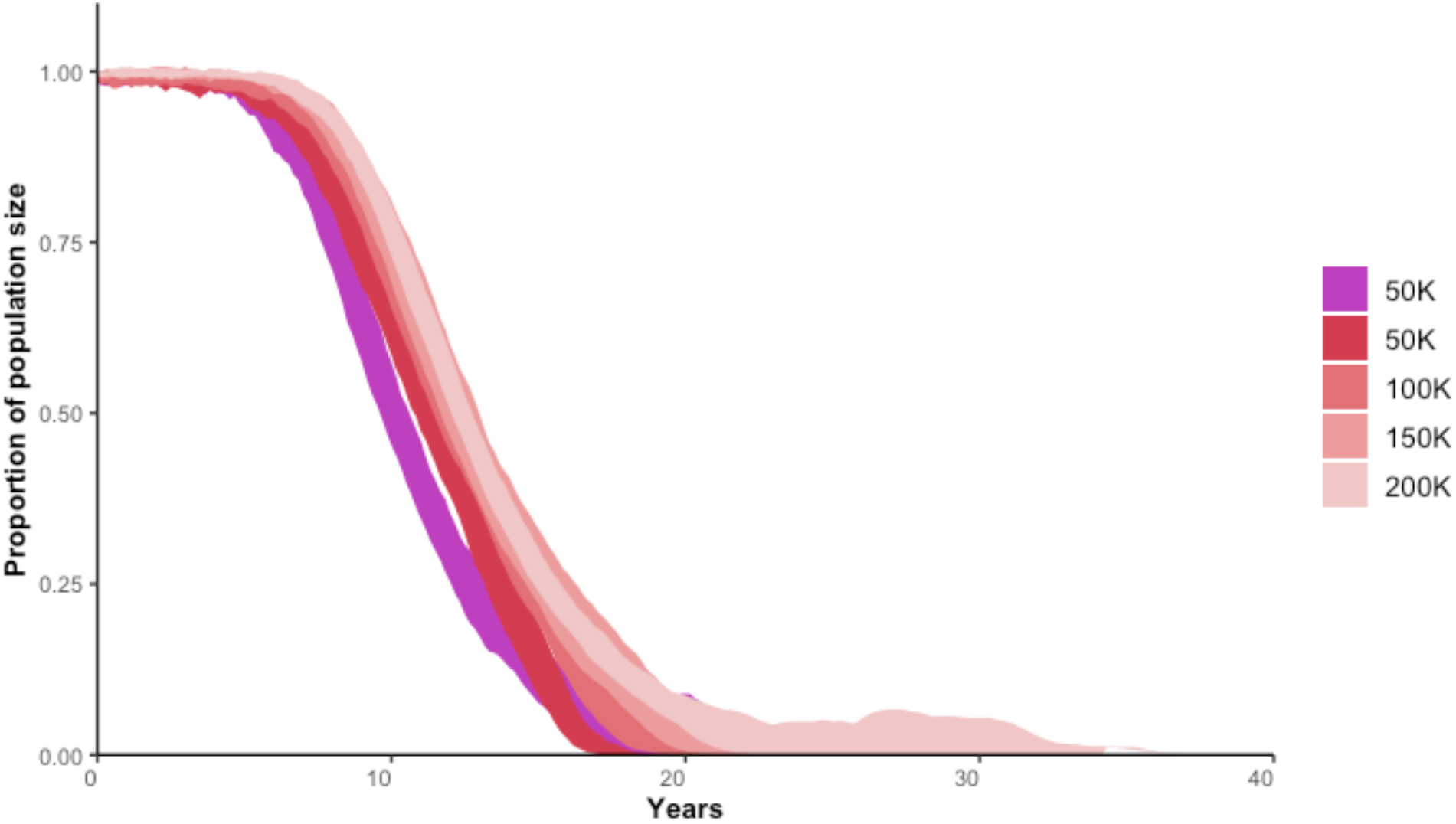
Proportion of population size after the inoculation with *t*_CRISPR_ individuals at year 0 based on 20 simulations where island size remained the same but the carrying capacity (hence the population density) were different (shades of red), and population density were the same but the island size changed (purple). The width of each line represents the minimum and the maximum value observed in the simulations at that time.

**Extended Data Figure 10.**
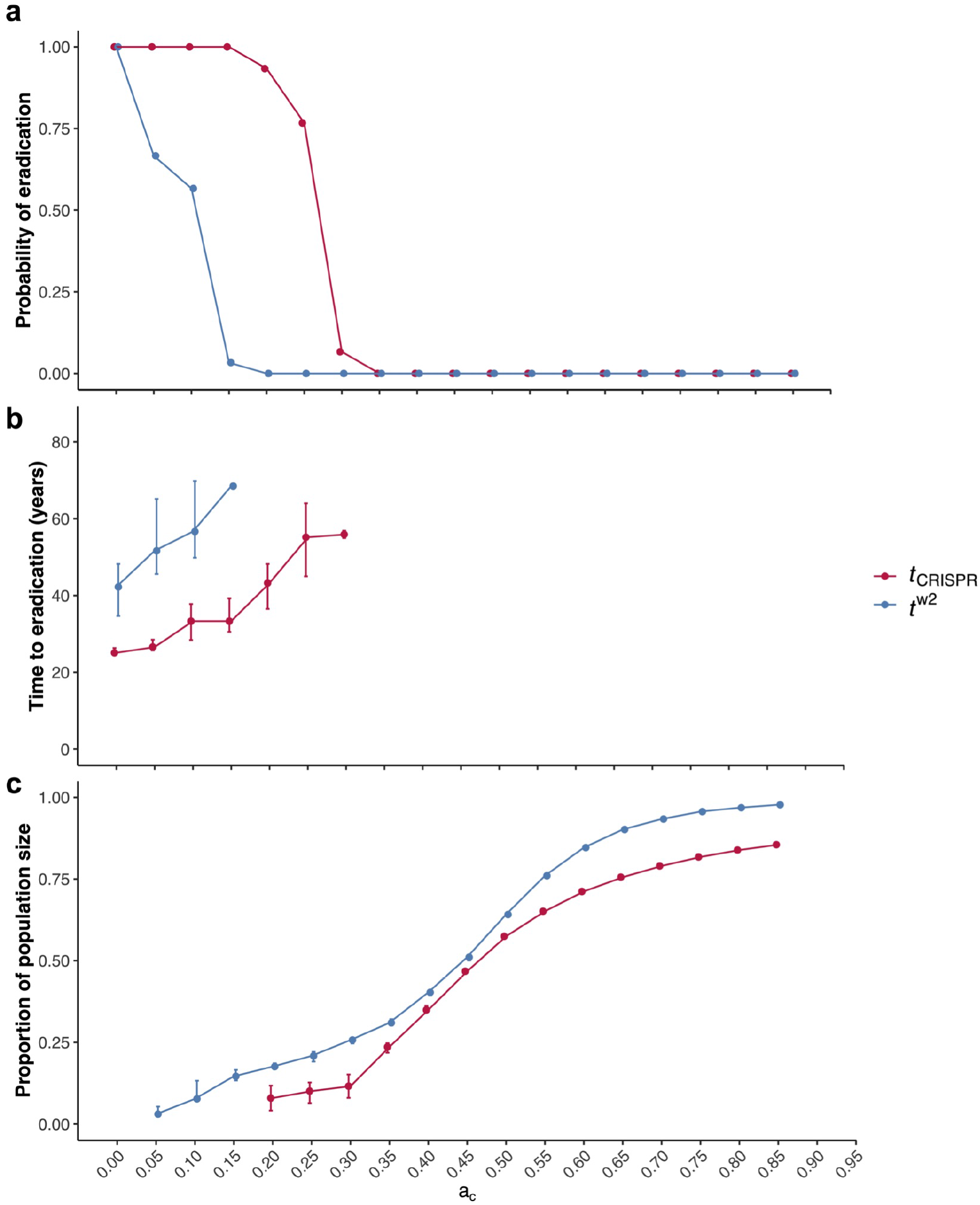
The influence of mate choice (*a*_c_ > 0) on **a.** the probability of eradication, **b**. the time to eradication, and **c**. the proportion of population size remaining when eradication was unsuccessful. Mate choice had an adverse effect on eradication; eradication failed when *a*_c_ > 0.15 using *t*^w2^, and when *a*_c_ > 0.30 using *t*_CRISPR_. Time to eradication was shorter with *t*_CRISPR_ than *t*^w2^, and *t*_CRISPR_ had better suppression when failed to eradicate (based on 30 simulations for each parameter combination, 1080 simulations in total).

**Extended Data Figure 11.**
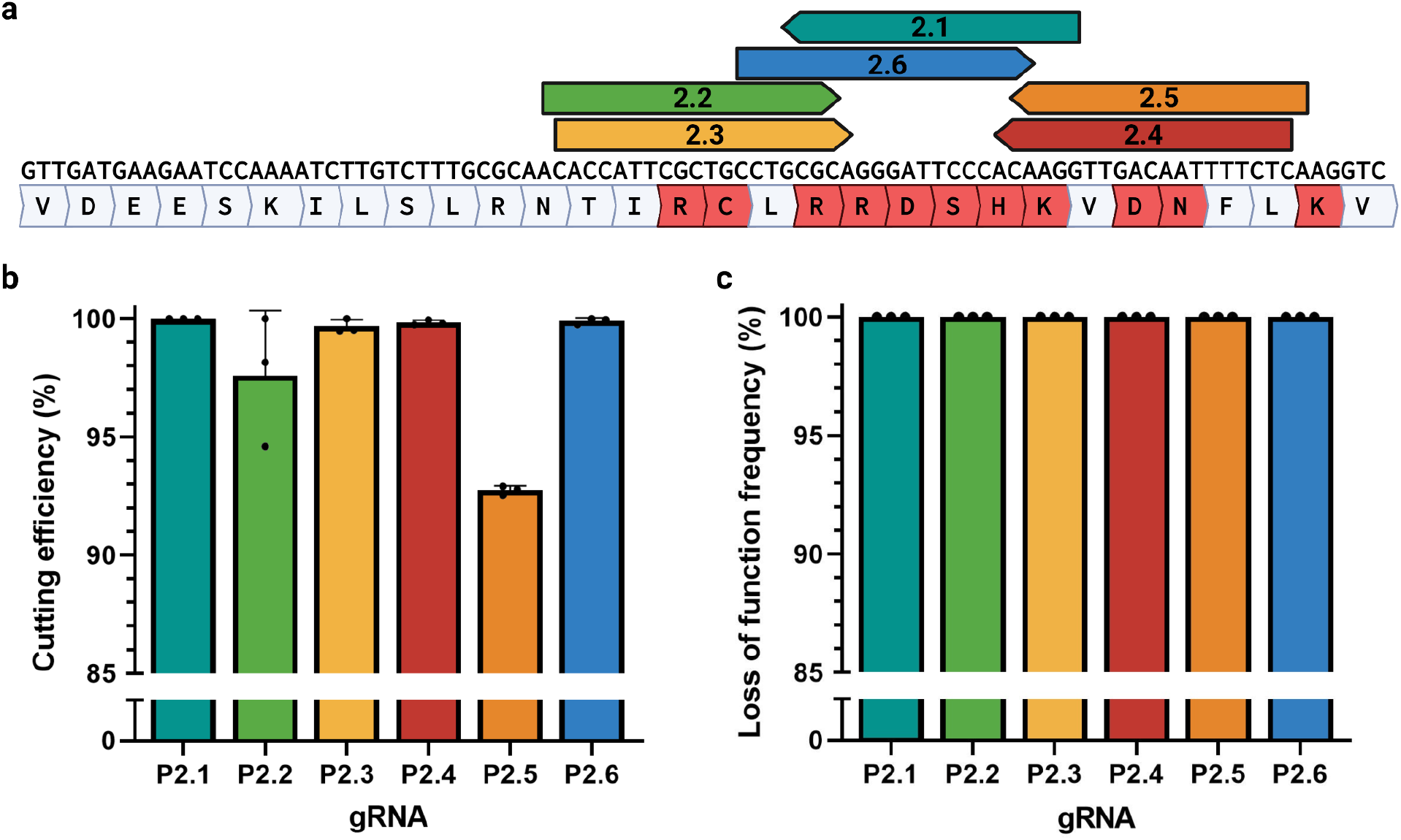
Selection and screening of *Prl* gRNAs. **a.** Position of gRNAs (2.1-2.6) targeting helix 4 of *Prl* (V159-V188). Amino acids in red have been demonstrated to have significant functional or structural roles and disruption of these is assumed to cause a loss of function. **b.c.** gRNAs were screened by transfecting mES cells followed by puromycin selection, DNA extraction and analysis by NGS. gRNA P2.3 was selected for further analysis due to its high cutting efficiency, loss of function frequency and ability to be combined with either P2.4 or P2.5. Bars show standard error of the mean.

**Extended Data Figure 12.**
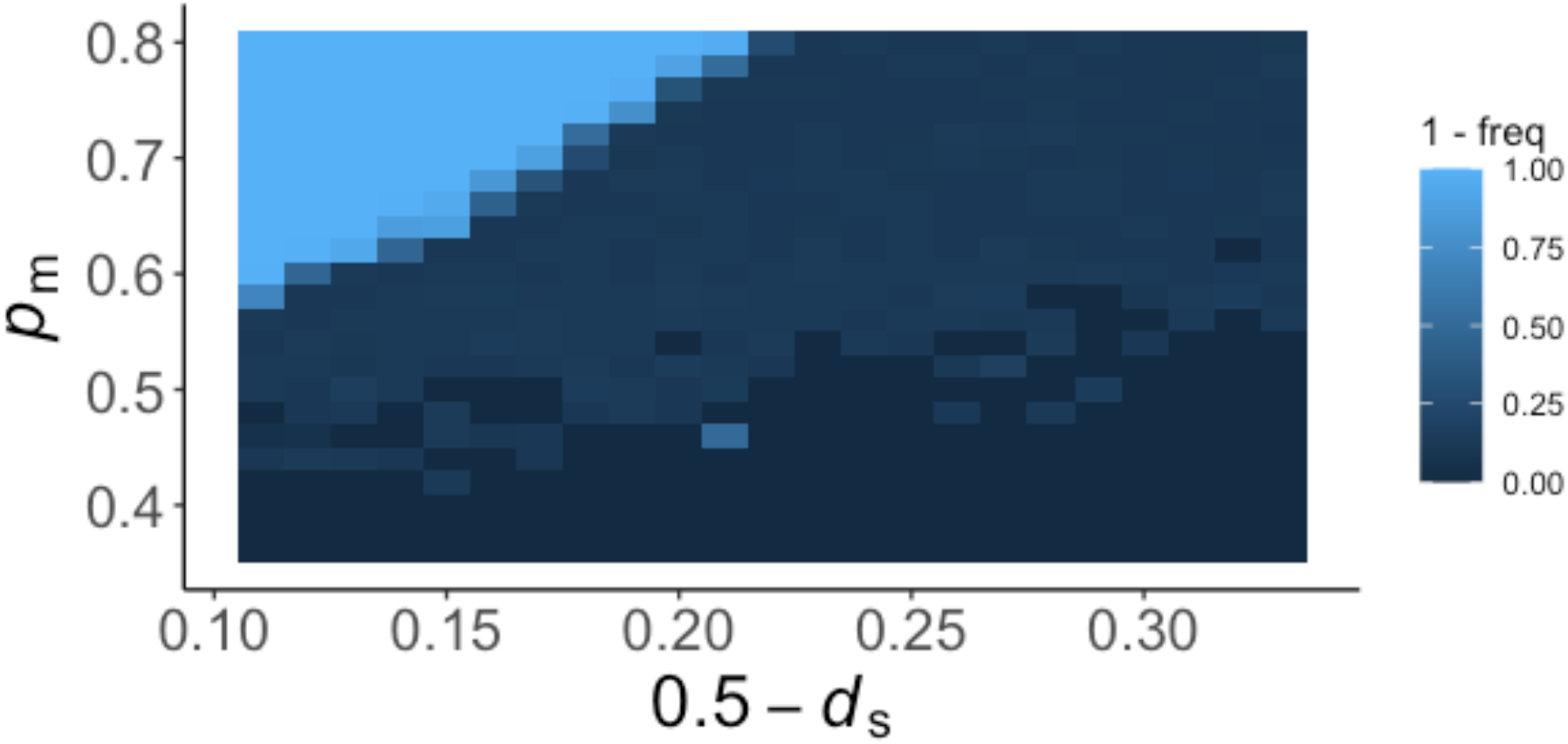
The frequency of *t*^w2^ at the end of simulations, which declines with increasing polyandry *p*_m_ and sperm competition *d*_s_ (based on one simulation for each parameter combination). Note that the x-axis and frequencies are displayed for easy comparison to Figure 3b in Manser *et al*. (2020)^31^.

**Extended Data Figure 13.**
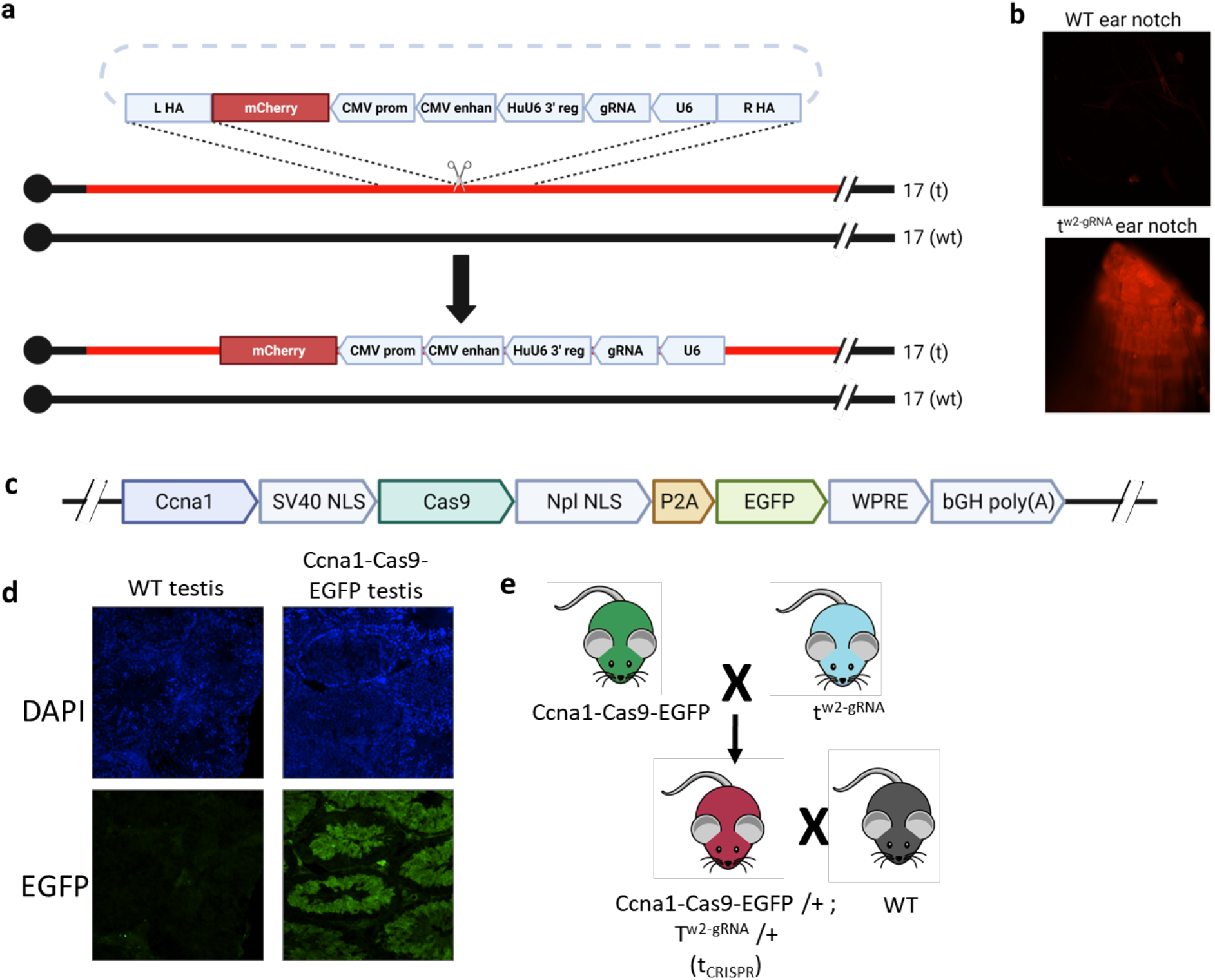
Generation and characterisation of split drive t_CRISPR_ mice. **a.b.**insertion of the gRNA and mCherry expression cassettes into the t haplotype using homologous recombination. *t*^w23-gRNA^ mice were genotyped by site specific PCR and mCherry fluorescence. **c.** Construct used to generate the Ccna1-Cas9-EGFP mice. **D.** DAPI (blue) and EGFP (green) immunofluorescence in adult testes correlated to Cas9 expression controlled by the Ccna1 promoter. **e.** Split drive breeding strategy to generate transhemizygous t_CRISPR_ mice.

## SUPPLEMENTARY INFORMATION

### *Prl* predicted loss of function allele analysis

Alleles were predicted to be loss of function if the reading frame was shifted or if any of the functionally important amino acid residues in helix 4^36–40^ (shown in red in Extended Data Figure 11) were missing (e.g. in frame deletion), substituted (e.g. point mutation) or positionally changed (e.g. upstream in frame deletion).

### Evidence of Cas9-gRNA carryover and activity in zygotes

Mosaic progeny can occur if CAS9 and gRNA are present in the gametes during fertilisation (commonly referred to as carry over). In insects, oocyte carryover has been observed at relatively high frequencies^92,93^ but is rare in sperm^93,94^ likely due to the difference in gamete size. Mosaic mice can have reduced fitness due to complete loss of function of the target gene in somatic cells reducing drive spread. Of the 142 offspring from transhemizygous *t*_CRISPR_ mice that contained indels, only one mosaic individual was detected indicating that sperm carryover is rare when using the *Ccna1* promoter and is not expected to impart a significant fitness cost.

